# PD-L1 blockade immunotherapy rewires cancer emergency myelopoiesis

**DOI:** 10.1101/2023.12.20.572561

**Authors:** Athina Boumpas, Antonis Papaioannou, Pavlos Bousounis, Maria Grigoriou, Veronica Bergo, Iosif Papafragkos, Athanasios Tasis, Michael Iskas, Louis Boon, Manousos Makridakis, Antonia Vlachou, Eleni Gavriilaki, Aikaterini Hatzioannou, Ioannis Mitroulis, Eirini Trompouki, Panayotis Verginis

**Author notes:** Panayotis Verginis, Laboratory of Immune Regulation and Tolerance, Division of Basic Sciences, Medical School, University of Crete, Heraklion 715 00, Greece.

## Abstract

Immune checkpoint blockade (ICB) immunotherapy has revolutionized cancer treatment, demonstrating exceptional clinical responses in a wide range of cancers. Despite the success, a significant proportion of patients still fail to respond, highlighting the existence of unappreciated mechanisms of immunotherapy resistance. Delineating such mechanisms is paramount to minimize immunotherapy failures and optimize the clinical benefit. Herein, we reveal that immunotherapy with PD-L1 blockage antibody (αPDL1) in tumour-bearing mice targets the hematopoietic stem and progenitor cells (HSPCs) in the bone marrow (ΒΜ), mediating their exit from quiescence and promoting their proliferation. Notably, disruption of the PDL1/PD1 axis induces transcriptomic reprogramming in HSPCs, from both individuals with Hodgkin lymphoma (HL) and tumour-bearing mice shifting towards an inflammatory state. Functionally, transplantation of HSPCs isolated from αPDL1-treated tumor-bearing mice exhibited resistance to cancer-associated myelopoiesis as evident by the generation of reduced frequencies of myeloid-derived suppressor cells (MDSCs) compared to cells from control-treated mice. Our findings shed light on unrecognized mechanisms of action of ICB immunotherapy in cancer, which involves targeting of BM-driven HSPCs and reprogramming of emergency myelopoiesis.

## Introduction

Immune checkpoint blockade (ICB) immunotherapy revolutionized cancer treatment, offering major therapeutic advantages across various cancer types as well as durable clinical responses in cancer patients (1, 2). As of to date, ICB immunotherapy targets the programmed cell death 1 (PD-1) and its ligand (PD-L1) (3), as well as cytotoxic T lymphocyte antigen 4 (CTLA-4) (4), which mediate dominant immunosuppressive signals. Despite the enormous success, the overall response rates to ICB immunotherapy remain low, highlighting the existence of unappreciated mechanisms of immune checkpoint resistance. Importantly, patients that respond to ICB often develop life-threatening immune-related adverse events (irAEs), which present a significant drawback in clinical applications (5, 6). There is an urgent need to comprehensively discern the mechanisms of action of ICB to optimize their therapeutic efficacy and minimize adverse events. Among the three ICB targets, the PD-L1 receptor has gained particular interest since it is broadly expressed by host and cancer cells. To this end, PD-L1 is expressed by myeloid cells such as dendritic cells (DCs), myeloid-derived suppressor cells (MDSCs) and macrophages, activated T and B lymphocytes, fibroblasts, and by certain epithelial cells upon inflammatory signaling (7). Notably, tumour cells also express various levels of PD-L1, which is considered as an immune escape mechanism (8). Although it is evident that the expression of PD-L1 by immune cells and tumour cells is required to promote tumour immune evasion and growth (9), still the precise mechanisms through which PD-L1 targeting contributes to the development of anti-tumour immunity remain poorly understood. Previous studies have shown that αPDL1 can reinvigorate exhausted CD8^+^ T cells (10) and facilitate the *de novo* priming of cytotoxic responses in the tumour-draining lymph nodes by interfering with the antigen-presentation capacity of DCs (11). Furthermore, PD-L1 blockade of human DCs induced the activation of caspase-1/ NLRP3-inflammasome and the release of inflammasome-dependent cytokines (12). Also, PD-L1 blockade triggered an inflammatory signature to mouse macrophages *in vivo* and *in vitro,* promoting anti-tumour immunity (13). Interestingly, treating asymptomatic multiple myeloma (AMM) patients with atezolizumab, induced an inflammatory signature in CD14^+^ monocytes (12), proposing that the PD-L1 axis may shape the myeloid-mediated inflammatory responses. In line with this hypothesis, myeloid skewing has been also implicated in resistance to αPDL1 immunotherapy. For example, the neutrophil-to-lymphocyte ratio (NLR) in the periphery of lymphoma patients has been proposed to correlate with the lack of response to αPDL1 treatment (14). Similarly, in advanced non-small cell lung cancer patients, responses to αPDL1 were associated with decreased frequencies of regulatory T cells (Tregs) (15, 16) and MDSCs as well as a reduction in NLR after treatment (16). However, whether αPDL1 immunotherapy imprints on the emergency myelopoiesis during cancer to alter the myeloid output in the periphery or directly modulates mature myeloid cell pool remains to be investigated. In addition, to what extent αPDL1 immunotherapy alters cancer-related myelopoiesis to promote cancer regression while fueling autoimmune adverse events remains elusive. Addressing these unanswered questions will provide important insights towards the design of rational therapies aiming to overcome αPDL1 resistance and to limit undesired systemic events.

Herein, we demonstrate that αPDL1 immunotherapy targets the hematopoietic stem and progenitor cells (HSPCs) in the bone marrow (BM), regulating the cancer-induced emergency myelopoiesis. Specifically, αPDL1 treatment increases the frequencies of BM-derived HSPCs and promotes their exit from quiescence in mice inoculated with either immunogenic or non-immunogenic tumours. Importantly, transcriptomic analysis showed that blocking the PDL1/PD1 axis induces inflammatory reprogramming in HSPCs from mice and individuals with Hodgkin lymphoma (HL). Additionally, transplantation of αPDL1-treated HPSCs significantly altered the cancer emergency myelopoiesis by reducing the frequencies of peripheral MDSCs. Overall, our findings shed light on unappreciated mechanisms of ICB immunotherapy, which consist of targeting the BM HSPCs and rewiring cancer emergency myelopoiesis, while providing new directions for understanding ICB immunotherapy resistance as well as the development of irAEs.

## Results

### Immunotherapy with αPDL1 contracts the MDSC compartment in tumour-bearing mice

To gain an in-depth understanding of the mechanisms underlying the responses to αPDL1 immunotherapy, we first analyzed the PD-L1 expression in major immune cell populations upon αPDL1-treatment (clone MIH5) of tumour-bearing mice. Flow cytometric analysis of mice bearing the non-immunogenic B16.F10-melanoma cell line demonstrated significantly decreased Geometric Mean Fluorescent Intensity (GMFI) of PD-L1 surface expression in CD3^+^ lymphocytes and CD11c^+^ DCs in both the tumour and spleen as well as on intratumoural CD11c^-^CD11b^+^Gr1^+^ MDSCs, when treated with αPDL1, compared to control (Fig. 1 A and B). In regard to frequencies, CD3^+^ T cells were not altered in the tumour nor the spleen (data not shown) of the control and αPDL1 treated melanoma-bearing mice. Frequencies of B16.F10 intratumoural DCs were also not affected by αPDL1, while MDSCs were significantly reduced (Fig. 1C and D, and EV1A). Similar results were observed in mice bearing the immunogenic MB49 bladder carcinoma cell line as well (Fig. 1E). Decreased frequency of MDSCs was also evident in the peripheral blood of αPDL1 treated B16.F10 bearing mice compared to control, while DC frequency remained unaltered (Fig. EV1B and C). In the spleen, both MDSCs and DCs decreased in the αPDL1-treated compared to control-treated animals bearing either melanoma (Fig. 1F and G, and EV1D) or bladder cancer (Fig. 1H). Further analysis of the splenic myeloid compartment showed that the frequencies of the CD11c^-^CD11b^high^Ly6C^+^Ly6G^-^ monocytic MDSC subset (M-MDSCs) significantly decreased in αPDL1-treated mice inoculated with B16.F10 (Fig. EV1E and F), yet in MB49-bearing presented no differences (Fig. EV1G), nor did the CD11c^-^CD11b^high^Ly6C^-^Ly6G^+^ granulocytic MDSC subset (G-MDSCs) in both tumour models (Fig. EV1E-G). Overall, these findings demonstrate a significant contraction of the MDSC compartment in the spleen, blood, and tumour of both immunogenic and non-immunogenic tumour-bearing animals upon αPDL1 treatment.

**Figure 1.**
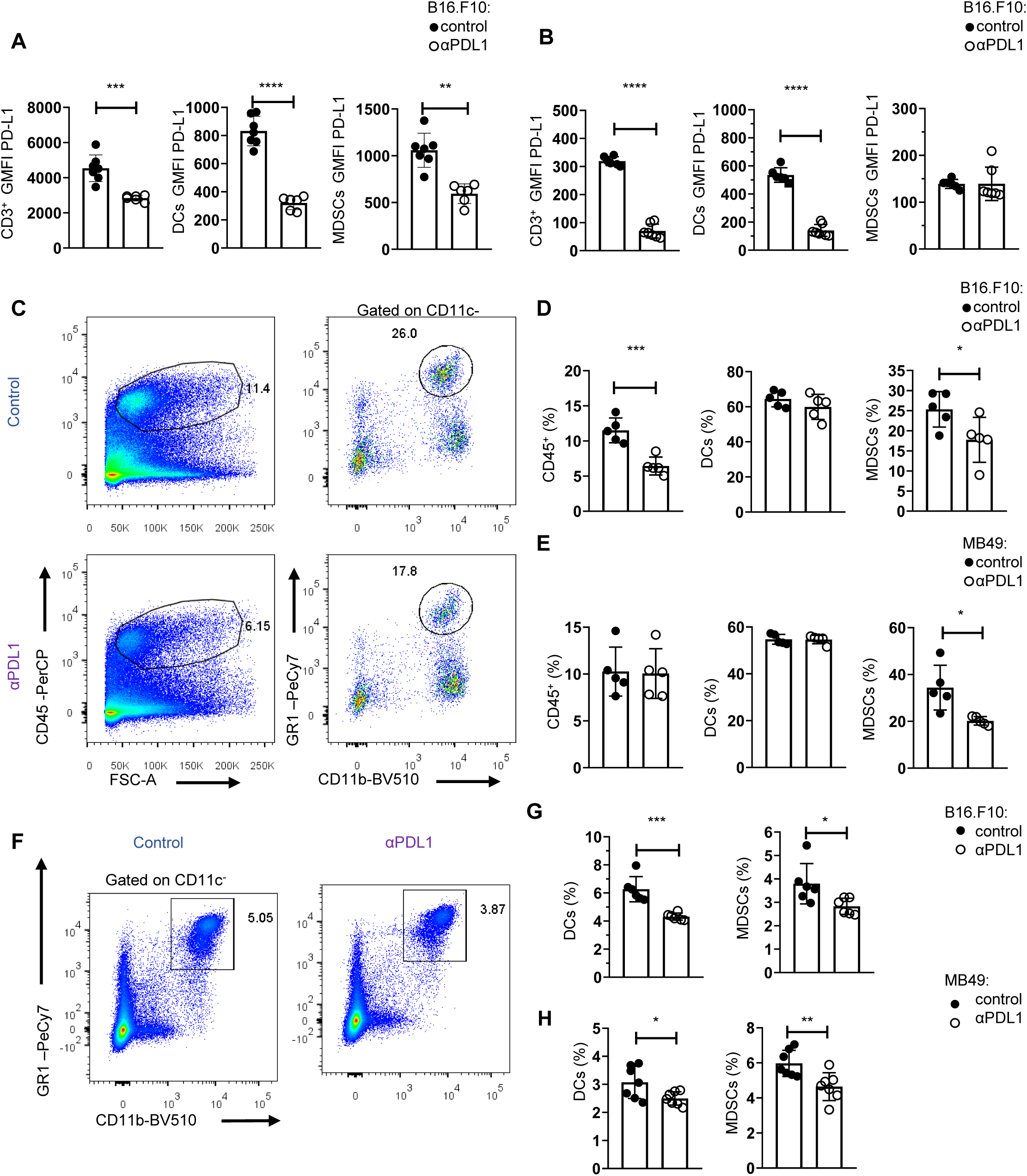
αPDL1 immunotherapy reduces peripheral MDSC frequencies during tumour progression. (**A**-**B**) Quantification through Flow Cytometry, of the GMFI of PD-L1 surface expression measurements of intratumoural (**A**; n = 7 control, n = 6 αPDL1) and splenic (**B**; n = 6 control, n = 7 αPDL1) CD3^+^ cells, CD11c^+^ DCs, and CD11c^-^CD11b^+^Gr1^+^ MDSCs in eight days B16.F10 melanoma bearing C57BL/6 mice treated either with PBS or αPDL1. Representative data from 4 independent experiments. (**C**-**E**) Representative FACs plots (**C**; Numbers denote percentages of gated populations) and quantification (**D**) of intratumoural CD45^+^ cells and MDSCs in PBS or αPDL1 treated C57BL/6 mice after eight days of B16.F10 (**C**, **D**; n = 5 control, n = 5 αPDL1) and twelve days of MB49 (**E**; n = 5 control, n = 5 αPDL1) tumour progression. Data from 1 experiment (**D**, **E**). (**F**-**H**) Representative FACS plots (**F**; Numbers denote percentages of gated populations) and frequencies of splenic MDSCs during the eighth day of B16.F10 (**F**, **G**; n = 6 control, n = 6 αPDL1) and MB49 (**H**; n = 7 control, n = 7 αPDL1) tumour progression in C57BL/6 mice treated with PBS or αPDL1. Data from 2 independent experiments (**G**), and data from two combined independent experiments (**H**). p < 0.05*, p < 0.01**, p < 0.001***, p < 0.0001****. If not stated otherwise, unpaired two-tailed t-tests are performed. Means and SEM are depicted in all bar plots. n = biologically independent mouse samples

To examine whether αPDL1 treatment could also imprint on the functional properties of MDSCs, we performed an *in vitro* suppression assay. To this end, M-MDSCs and G-MDSCs isolated from the spleen of control and αPDL1-treated melanoma-bearing mice and were co-cultured with CellTrace Violet (CTV) labelled T effector (CD4^+^Foxp3^-^) cells sorted from naïve Foxp3^EGFP^ mice in the presence of anti-CD3/anti-CD28 activation beads (Fig. EV2A). Both MDSC subsets from αPDL1 treated mice displayed a sustained suppressive ability compared with the control group, as evident by the similar CTV dilution (Fig. EV2B), and decreased T cell activation based on the CD44 and CD25 expression (data not shown). Collectively, these results demonstrate that αPDL1 treatment significantly decreases the frequencies of MDSCs in tumour-bearing mice without altering their functional properties.

### Human and mouse BM HSPCs express PD-L1

Since αPDL1 treatment contracts the MDSC compartment in the periphery of tumour-bearing animals and considering that MDSCs originate from ΒΜ hematopoietic progenitors (17), we hypothesized that αPDL1 may regulate cancer emergency myelopoiesis. To address this, we first examined whether HSPCs express PD-L1, and if they constitute a target of αPDL1 immunotherapy. To this end, Flow cytometric analysis showed that HSPCs (or LSK cells; Lineage (Lin)^-^Sca1^+^cKit^+^) express PD-L1 at steady state and its expression was not altered after B16.F10 melanoma cell inoculation (Fig. 2A), whereas it was significantly increased upon inoculation with immunogenic MB49 bladder cancer cells (Fig. 2B). In support, BM stem (CD45^low^CD34^+^CD38^-^, Fig. 2C and D) and progenitor (CD45^low^CD34^+^CD38^+^, Fig. 2C and E) cells from patients with HL express PD-L1 as compared to fluorescence minus one (FMO) staining.

**Figure 2.**
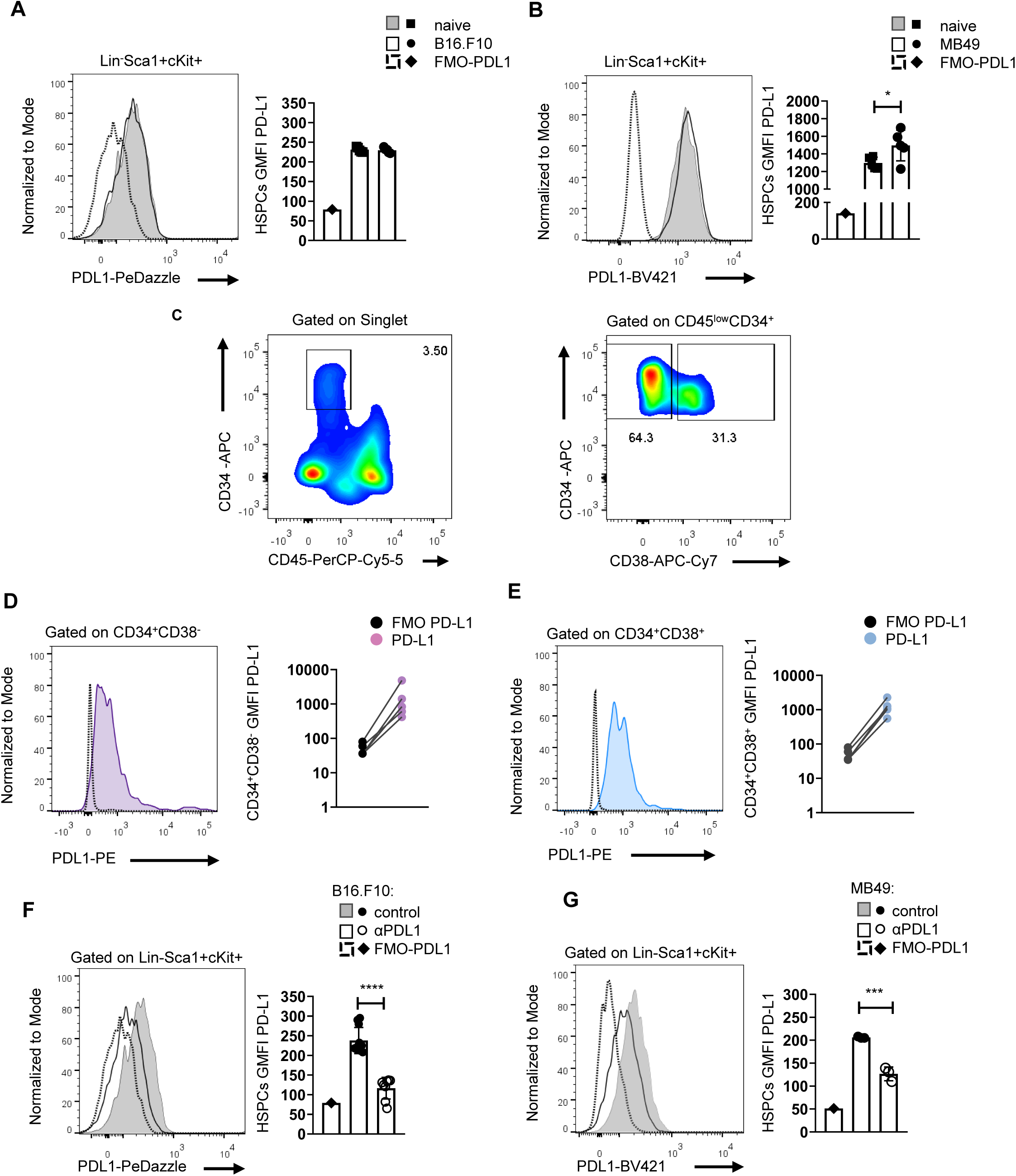
Murine and human HSPCs express PD-L1. (**A**-**B**) Representative histograms (left) and quantification (right) of surface PD-L1 expression in murine BM HSPCs eight days following B16.F10 (**A**; n = 5 naïve, n = 5 B16.F10) or MB49 (**B**; n = 5 naïve, n = 5 MB49) inoculation in C57BL/6 mice. Representative Flow cytometry data from 1 (**B**), and 2 (**A**) independent experiments. (**C**-**E**) Representative gating strategy (**C**; Numbers denote percentages of gated populations) of human BM stem (CD45^low^CD34^+^CD38^-^), and progenitors (CD45^low^CD34^+^CD38^+^) cells isolated from HL patients isolated at diagnosis. Representative overlays (left) and GMFI quantification (right) of PD-L1 in CD34^+^CD38^-^ (**D**; n = 5) and CD34^+^CD38^+^ (**E**; n = 5) HL patients compared to their counterpart FMO (representation in a Log10 scale). (**F**, **G**) Representative histograms (left) and quantification (right) of PD-L1 surface expression in BM HSPCs during the eighth day of B16.F10 (**F**; n = 8 control, n = 11 αPDL1) or MB49 (**G**; n = 3 control, n = 3 αPDL1) tumour development in C57BL/6 mice treated with PBS or αPDL1. Representative Flow cytometry data from 2 (**G**), and 10 (**F**; 2 of them are displayed) independent experiments. p < 0.05*, p < 0.01**, p < 0.001***, p < 0.0001****. If not stated otherwise, unpaired two-tailed t-tests are performed. Means and SEM are depicted in all bar plots. n = biologically independent mouse or human samples

Notably, treatment with αPDL1 (clone MIH5) resulted in significantly decreased staining of PD-L1 (clone MIH5 or 10F.9G2) in HSPCs of mice inoculated with either non-immunogenic (Fig. 2F) or immunogenic tumour cells (Fig. 2G), pointed to specific targeting of the HSPC compartment. Together, these results establish that HSPCs in the BM express PD-L1 and are targeted by αPDL1 immunotherapy in tumour-bearing mice.

### αPDL1 treatment induces the expansion of HSPCs and their exit from quiescence

We next asked whether αPDL1 targeting the HSPCs in the BM affects myelopoiesis. To address this, we first examined if αPDL1 treatments affect the frequencies of HSPCs during cancer. Notably, αPDL1 treatment significantly expanded the HSPC compartment in mice inoculated with either non-immunogenic (Fig. 3A and B, and EV3A) or immunogenic tumour cells (Fig. 3C and D) compared to control-treated mice. Furthermore, the frequencies of CD48^−^CD150^+^ long-term HSCs (LT-HSCs) were not altered, CD48^−^CD150^-^ short-term HSCs (ST-HSCs) significantly decreased, and multipotent progenitors CD48^+^CD150^-^ (MPPs) were significantly increased in αPDL1-treated B16.F10-injected (Fig. 3B) and MB49-injected (Fig. 3D) mice. These results demonstrate that αPDL1 treatment expands the HSPC compartment in the BM of tumour-inoculated animals.

**Figure 3.**
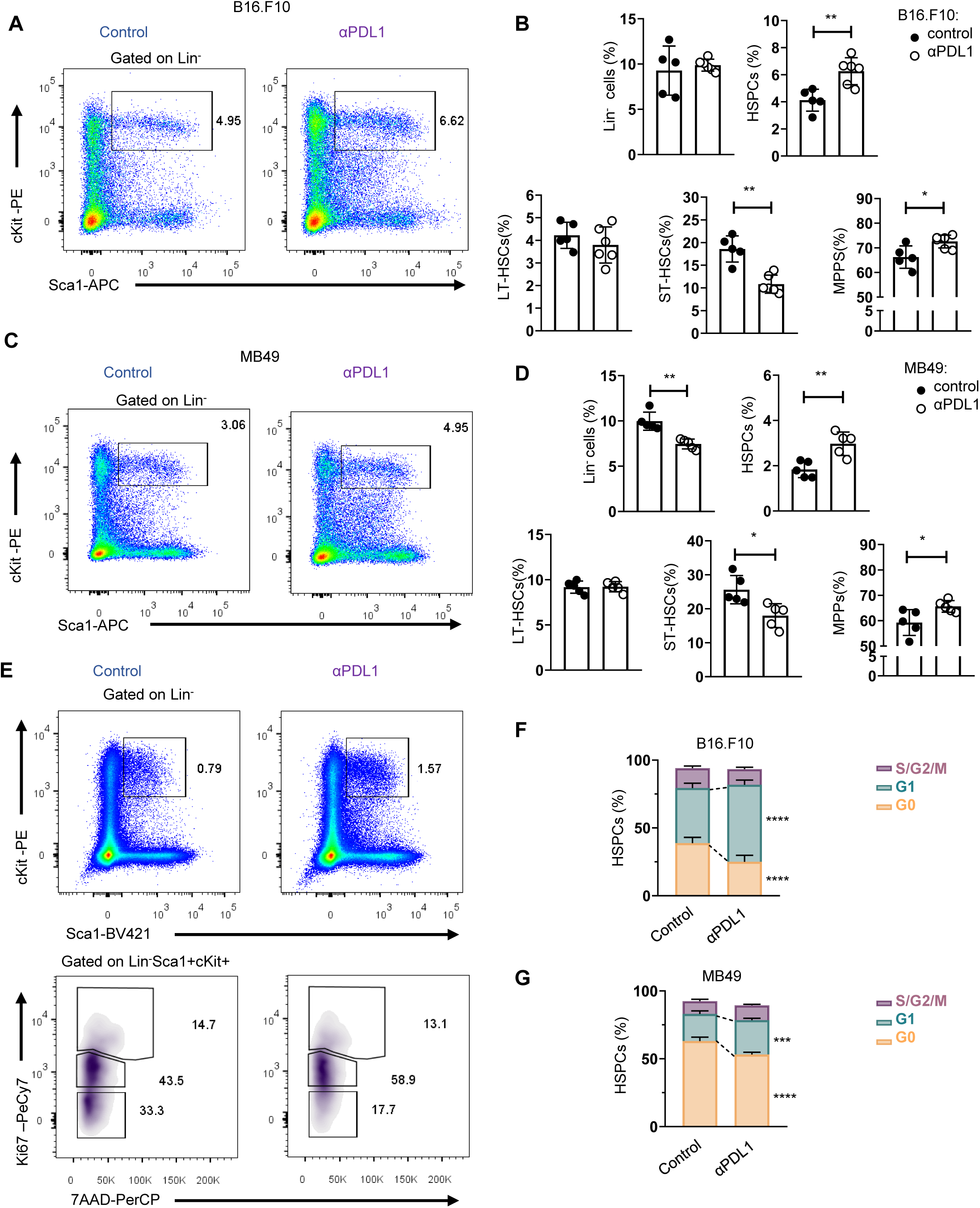
Administration of αPDL1 drives expansion of the HSPC compartment and promotes their activation. (**A**-**D**) Representative FACs plots (**A, C**; Numbers denote percentages of gated populations) and the percentages of BM HSPCs (LSK: (Lin)^-^Sca1^+^cKit^+^), including the three subpopulations categorized according to their CD150 and CD48 expression pattern, LT-HSCs, ST-HSCs, and MPPs in PBS or αPDL1treated C57BL/6 mice inoculated with B16.F10 (**B**; n = 6 control, n = 6 αPDL1) and MB49 (**D**; n = 3 control, n = 3 αPDL1) and sacrificed after eight days. Representative data from 1 (**D**; HSPCs subsets), 2 (**D**; HSPCs), 3 (**B**; HSPC subsets), and 8 independent experiments (**B**; HSPCs). (**E**-**F**) Representative FACs plots (**E**; Numbers denote percentages of gated populations) of BM HSPCs isolated from PBS- or αPDL1-treated C57BL/6 mice inoculated with B16.F10 (**E**-**F**; n = 5 control, αPDL1 n = 5) and MB49 (**G**; n = 4 control, n = 5 αPDL1). After eight days mice sacrificed and stained with Ki67 and 7-AAD for cell cycle analyses. Frequencies of HSPCs (**F**, **G**) in the G0, G1, and S/G2/M cell cycle phases. Two-way ANOVA was performed. Data from 1 experiment (**F**, **G**). p < 0.05*, p < 0.01**, p < 0.001***, p < 0.0001****. If not stated otherwise, unpaired two-tailed t-tests are performed. Means and SEM are depicted in all bar plots. n = biologically independent mouse samples

To examine whether αPDL1 treatment actively induces HSPC proliferation, we performed flow cytometry analysis upon staining with 7-AAD to measure cell DNA content and Ki-67 to distinguish proliferating from non-proliferating/quiescent cells. Indeed, αPDL1-treated B16.F10-(Fig. 3E and F) and MB49-inoculated (Fig. 3G) animals showed reduced percentages of HSPCs in the G0 phase and significantly increased percentages in the G1 phase compared to control-treated mice. Taken together, these findings provide evidence that αPDL1 immunotherapy promotes the exit of HSPCs from the quiescent state and induces their proliferation in the BM.

We next asked if αPDL1 alters the HSPCs differentiation potential during the early stages of myeloid commitment. Therefore, we assessed frequencies of committed myeloid progenitors (LK: Lin^-^Sca1^-^cKit^+^) that can further differentiate into common myeloid progenitors (CMP; LK CD34^+^CD16/32^−^), granulocyte-macrophage progenitors (GMPs; LK CD34^+^CD16/32^+^) and megakaryocyte-erythrocyte progenitors (MEPs; LK CD34^−^CD16/32^−^). Interestingly, αPDL1 treatment of non-immunogenic tumour-bearing mice did not affect the frequencies of the myeloid progenitors (Fig. EV4A and B) but the frequency of common lymphoid progenitors (CLPs: Lin^-^Sca1^low^cKit^low^IL7Ra^high^CD135^high^) was significantly increased (Fig. EV4D and E). Contrarily, αPDL1 treatment decreased the frequency of CMPs in immunogenic tumours, while increasing the frequency of GMP (Fig. EV4C) without affecting the CLP frequency (Fig. EV4F). Collectively, these findings suggest that αPDL1 treatment imprints on the expansion of the HSPC compartment, while the immunogenicity of the tumour dictates its differentiation potential.

### Targeting the PD1/PDL1 axis expands the HSPC compartment in the BM

To examine whether the expansion of HSPCs in tumour-bearing mice is specific to αPDL1, we treated mice with either αPD1 or αCTLA4, the two ICBs used in the treatment of patients with solid malignancies. Interestingly, only αPD1 treatment of B16.F10-injected mice demonstrated a significant increase of HSPCs frequency, whereas no difference was observed in αCTLA4-treated mice (Fig. 4A and B). These results suggest that interfering with specifically the PD1/PDL1 axis promotes the expansion of the HSPC compartment. Additionally, a significant increase in HSPCs frequencies was observed upon B16.F10 cell inoculation of PD1-deficient (*PD1^-/-^*) compared to WT animals (Fig. 4C). Although PD-1 has been shown to be expressed by various cell types of hematopoietic origin (18, 19), T cells constitute the major source of PD-1 expression (18, 20). Therefore, to provide mechanistic insights into our findings, we asked whether PD-1 expression by T-cells contributes to the expansion of the HSPC compartment in αPDL1-treated animals. To this end, *RAG1^-/-^* immunodeficient animals, which do not harbor T or B lymphocytes due to a defect in the receptor recombination mechanism (21), were inoculated with B16.F10 cells and treated with αPDL1. Surprisingly, although αPDL1 efficiently targeted the HSPCs (Fig. 4E), no significant differences were observed in their frequencies between αPDL1- and control-treated *RAG1^-/-^* mice (Fig. 4D and F). Collectively, these results suggest that targeting the PD1/PDL1 axis mediates the HSPC compartment’s expansion during cancer development and highlights an essential role of lymphocytes in this process.

**Figure 4.**
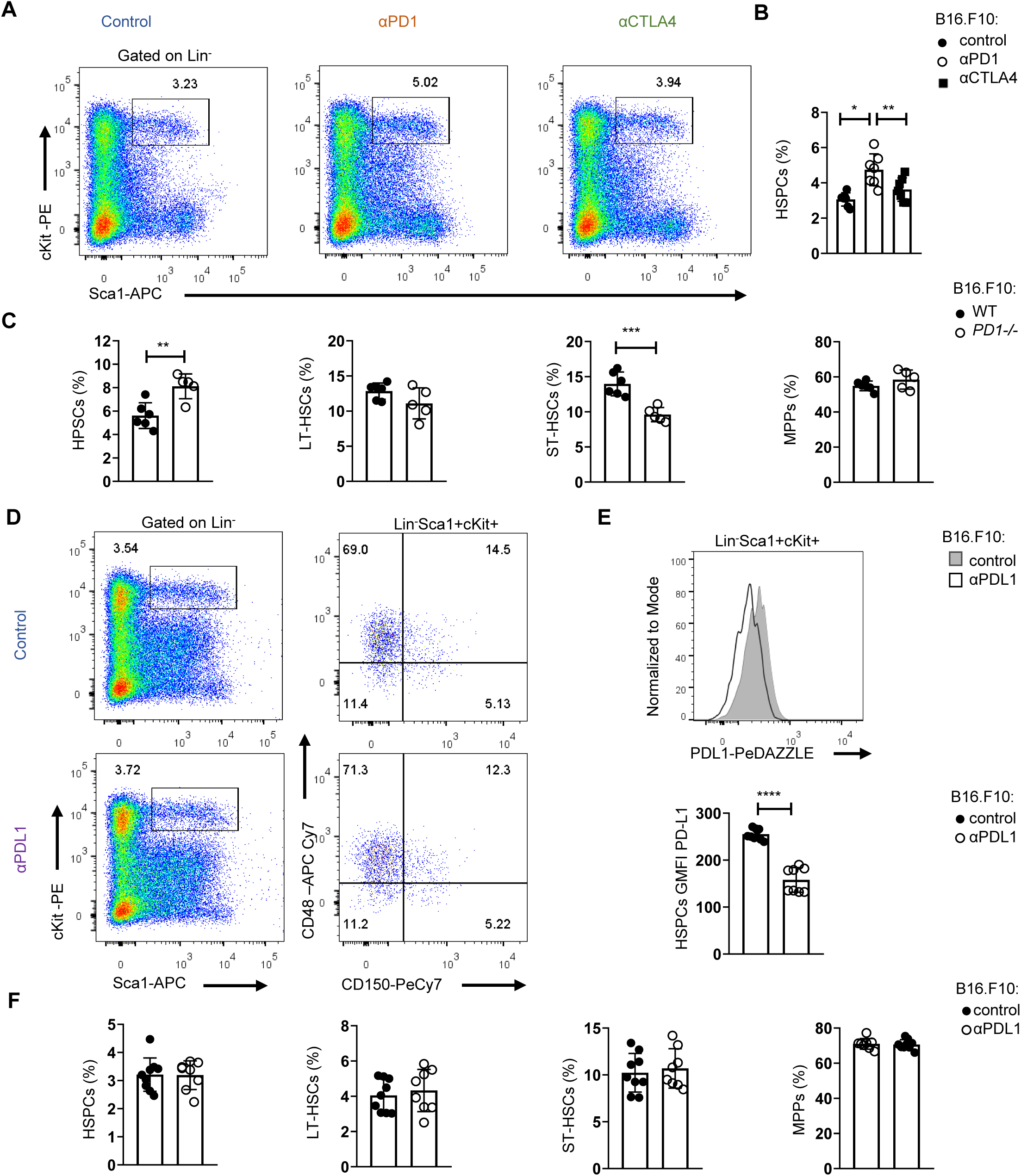
Targeting of the PDL1/PD1 axis expands the HSPC compartment in the BM. (**A**-**B**) Representative FACS plots (**A**; Numbers denote percentages of gated populations) and HSPCs frequencies (**B**; n = 7 control, n = 7 αPD1, n = 7 αCTLA4) in C57BL/6 mice inoculated with B16.F10 and treated with either αPD1, αCTLA4, or PBS with BM analysis performed after eight days. Data from 2 combined independent experiments. One-way ANOVA was performed. (**C**) Frequencies of BM HSPCs, and HSPCs subsets isolated from B16.F10 inoculated *WT* or *PD1^-/-^* C57BL/6 mice (n = 6 WT, n = 5 *PD1*^-/-^), sacrificed during the eighth day of tumour development. Data from 1 experiment, assessed with Flow cytometry. (**D**-**E**) Representative FACs plots (**D**; Numbers denote percentages of gated population) and frequencies (**F**; n = 9 control, n = 8 αPDL1) of HSPCs and the HSPCs subpopulations in *RAG1^-/-^* mice inoculated with B16.F10 and treated with either αPDL1 or PBS with BM analysis performed after eight days. Representative histograms (up) and GMFI quantification (down) (**E** n = 9 control, n = 8 αPDL1) of the PD-L1 surface expression of the aforementioned HSPCs. Data from 2 combined independent experiments, assessed with Flow cytometry. p < 0.05*, p < 0.01**, p < 0.001***, p < 0.0001****. If not stated otherwise, unpaired two-tailed t-tests are performed. Means and SEM are depicted in all bar plots. n = biologically independent mouse samples

### Enhanced inflammatory signaling and altered myelopoiesis in HSPCs upon αPDL1 immunotherapy

To gain insights into the molecular mechanisms underlying the αPDL1-mediated expansion and differentiation of HSPCs in tumour-bearing animals, we first evaluated the proteome of BM sera from αPDL1-treated and control-treated tumour-inoculated mice. Seventy-nine (79) differentially expressed proteins (DEPs, p < 0.05) (Fig. EV5A) were identified, and forty-one (41) exhibited Fold Change |FC| ≥ 1.5 (seven (7), upregulated, thirty-four (34) downregulated). To this end, proteins related to hematopoiesis (i.e. *Kars*, *Serpina1c*), stress (*Stip1*, *Stk4*, *Gmps*) and inflammation (*Cndp2*, *Map2k1*, *Cfp*, *Sik2*) were significantly upregulated in sera from αPDL1-treated compared to control mice. Interestingly, the innate immune receptor melanoma differentiation-associated protein 5 (MDA5; encoded by *Ifih1*), which drives hematopoietic regeneration (22), was exclusively present in the sera from αPDL1-treated mice, while TGF-β signaling which has been linked to HSC quiescence (23), was downregulated in αPDL1-treated mice. In support, Gene Ontology analysis (GO) (Fig. EV5B), Gene Set Enrichment Analysis (GSEA) (Fig. EV5C), and Ingenuity Pathways Analysis (IPA) (Fig. EV5D), pointed to enhanced inflammation-induced pathways (GO: “TNF signaling pathway”, “Interleukin-1 family signaling”, IPA: “LXR/RXR activation”, “NRF2 mediated Oxidative stress response”, GSEA: “Inflammatory Response”), cell cycle (GO: “regulation of cell cycle G1/S phase”, IPA: “HIF1a signaling pathway”) and metabolic reprogramming (GO: “metabolism of nucleotides”, IPA: “Glycolysis”, “Integrins”, “iron homeostasis signaling pathway”) in αPDL1-treated group of mice. Transcriptomic analysis of HSPCs isolated from B16.F10 melanoma-bearing mice either αPDL1- or control-treated, revealed fifty-eight (58) differentially expressed genes (DEGs) (forty-five (45) upregulated, thirteen (13) downregulated, |FC| ≥ 1.5, FDR<0.05) (Fig. 5A). Among these, genes associated with stress response and inflammation (*Dusp1, Fos, Zfp36, Hspa5, Ier2*) were significantly upregulated in HSPCs from αPDL1-treated compared to control mice. Importantly, genes that regulate HSPC proliferation (*Klf4, Pf4, Cd69, Egr1*) and differentiation (*Klf2, Fosb, Jun, Klf6*) were also upregulated in αPDL-1-treated. This was also evident upon pathway analysis of DEGs which showed enrichment in “response to stress pathways”, “response to Cytokine” and “inflammation” (Fig. 5B). Supporting these results, GSEA pointed to positive enrichment “Negative Regulation Of Myeloid Cell Differentiation” (NES 1.44, FDR 0.18), “Hematopoietic Stem Cell Differentiation”, and “TNF-a Signaling via NF-kB” (Fig. 5C) of αPDL1 treated HSPCs compared to control. Finally, through IPA, inflammatory-related biological functions such as “S100 signaling”, “dendritic cell maturation” and “alternative macrophage activation” were predicted to be more active in HSPCs from αPDL1-treated tumour-bearing animals (Fig. 5D). Furthermore, transcriptomic analysis on HSPCs isolated from αPDL1-treated and control MB49-inoculated animals (|FC|>1.5, FDR<0.05) revealed seventy-six (76) DEGs (twenty-nine (29) upregulated, forty-seven (47) downregulated) (Fig. EV5E). Interestingly, genes associated with neutrophil development (*Camp, Mmp9, Ltf, Lrg1*) and maturation (*S100A, S100B, Retnlg, Cd177, Wfdc21*), myeloid differentiation (*Clec5a, Lcn2, Ngp, Lmna*) and monocyte activity (*Irf2bp2*) were significantly downregulated in αPDL1-treated group, whereas genes involved in cell self-renewal (*Egln1, Thbs1*) were significantly upregulated. GO analysis showed significant enrichment of terms such as “response to stress”, “myeloid differentiation” and “inflammatory response” in HSPCs isolated from αPDL1-treated mice compared to control animals (Fig. EV5F). Additionally, GSEA analysis indicated a negative enrichment in “myeloid cell differentiation” and positive enrichment of “Interferon alpha response” (Fig. EV5G). Collectively, these results demonstrate that αPDL1 immunotherapy causes transcriptomic reprogramming in HSPCs in the BM of both immunogenic and non-immunogenic cancer cell-inoculated animals.

**Figure 5.**
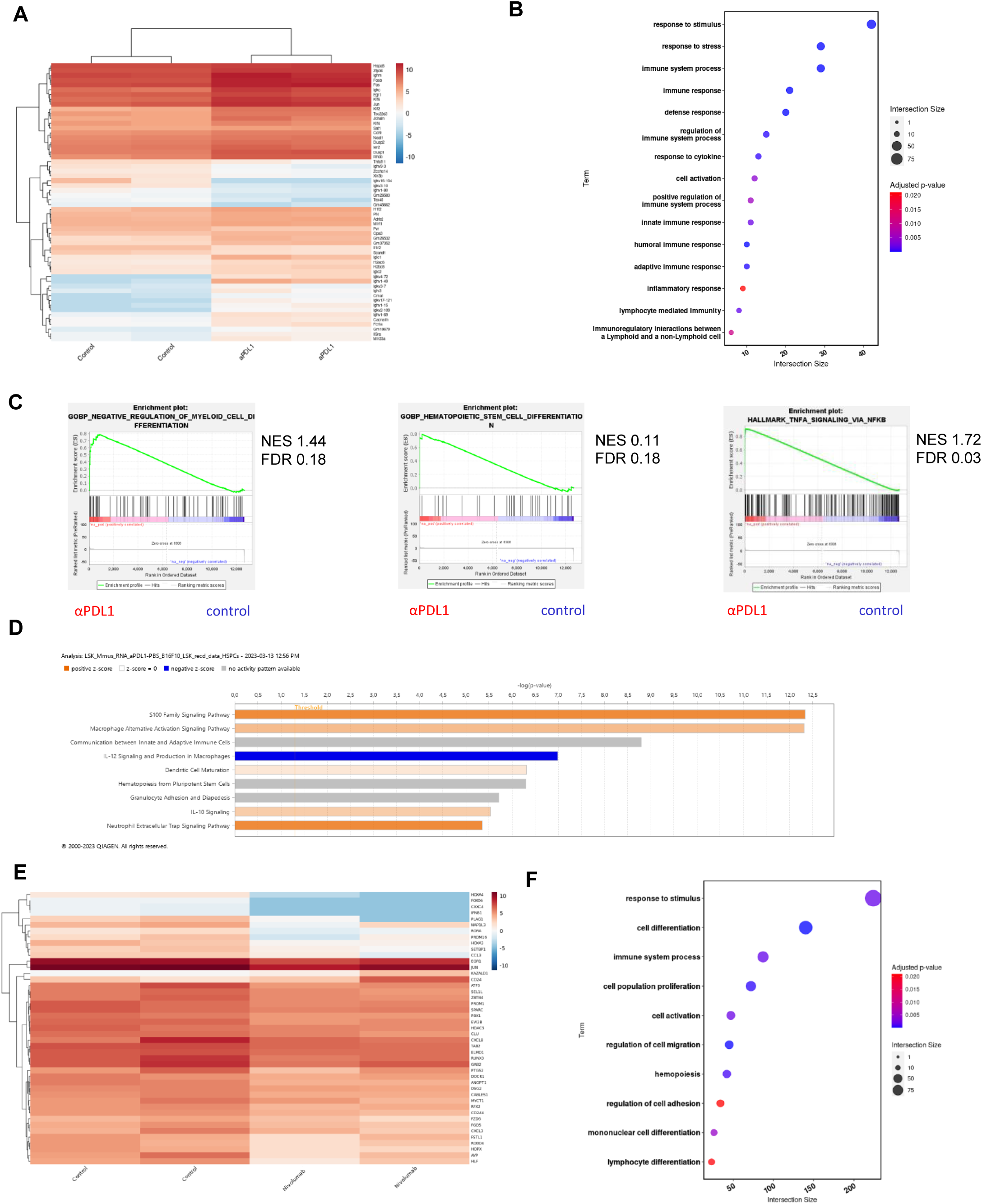
Immunotherapy induces transcriptomic reprogramming of BM HSPCs. (**A**) RNA-seq heatmap of the 58 DEGs (45 up-regulated and 13 down-regulated genes versus controls, |FC| ≥ 1.5, FDR < 0.05) of BM HSPCs isolated from B16.F10 tumour-bearing C57BL/6 mice treated with PBS (Control; n = 2) or αPDL1 (n = 2). (**G**) Pathway analysis of DEGs from BM HSPCs isolated from B16.F10 tumour-bearing C57BL/6 mice treated with PBS (n = 2) or αPDL1 (n = 2). (**C**) GSEA plot showing the positively enriched pathways “Negative Regulation Of Myeloid Cell Differentiation” (NES 1.44, FDR 0.18), “Hematopoietic Stem Cell Differentiation” (NES 0.11, FDR 0.18), “TNF-a signaling via NF-kB” (NES 1.72, FDR 0.03) of αPDL1 group compared to control (FDR (q-value) < 25%). (**D**) BM specific IPA, of signaling pathways in BM-HSPCs from melanoma αPDL1-reated mice, as compared to PBS-treated mice (control). Bar color reflects the IPA activation z-score of an enriched pathway which indicates the direction of effect associated from gene to pathway, with orange representing a direct, and blue representing an indirect association between pathway activation/inhibition and gene expression.(**E**) RNA-seq heatmap of representative DEGs of CD34^+^ cells from BM of HL patients isolated at diagnosis (n = 2) and αPD1-treated (Nivolumab; n = 2) HL patients (612 DEGs with 202 up-regulated and 410 down-regulated versus controls, FDR < 0.05). (**F**) Pathway analysis of DEGs of CD34^+^ BM cells isolated from untreated and αPD1 treated HL patients. n = biologically independent mouse and human samples

Interestingly, transcriptomic analysis of CD34^+^ cells from BM of αPD1-treated individuals with HL revealed six hundred and twelve (612) DEGs, of which two hundred and two (202) were upregulated and four hundred and ten (410) were downregulated (FDR< 0.05) compared to CD34^+^ cells from samples from HL patients isolated at diagnosis (Fig. 5E). Specifically, genes associated with HSC expansion (*CXCL8, DSG2, ZBTΒ4, MYCΤ1, PROM1, DOCK1*), self-renewal (*RORA, PLAG1, SPARC, SEL1L, PRDM16, PBX1, GAB2, CABLES1)*, proliferation (*EGR1, KAZALD1, ATF3*) and differentiation (*JUN, COL5A1*) were significantly downregulated in CD34^+^ cells from untreated compared to αPD1 treated individuals. Notably, genes that participate in HSC differentiation towards the myeloid cell lineage (*HOXA3, NAP1L3 RUNX3, RGS18, EVI2B, FZD6, CCL3)* were also downregulated in control group, while upregulation of *POU2AF1, RAG2, CD19, CD79A, CD79B* pointed to skewing towards development of lymphoid progenitors was evident in CD34^+^ cells form αPD1-treated individuals. GO further supported these findings with lymphocyte and myeloid differentiation pathways to be highly enriched as well as the response to stimulus pathway in accordance with the αPDL1-treated mouse data (Fig. 5F). Overall, the findings presented here, show that αPDL1 promotes a transcriptomic reprogramming of HSPCs underlined by inflammatory-related processes.

### αPDL1 immunotherapy modulates the myelopoiesis potential of HSPCs during cancer

We showed that αPDL1 immunotherapy promotes the exit of the HSPC compartment from quiescence and induce their transcriptomic re-wiring, raising the possibility of altered cancer myelopoiesis. To examine this hypothesis, we initially performed a Colony Forming Unit (CFU) assay, using medium formulated to optimally support the growth of primitive myeloid and erythroid progenitors of BM HSPCs isolated from αPDL1 or control-treated B16.F10 tumour-inoculated mice. CFU-monocytic (CFU-M) progenitors increased in αPDL1 cultured HSPCs, accompanied by a decrease of CFU-granulocytic (CFU-G), suggesting that αPDL1 treatment reprograms the myelopoiesis potential of HSPCs (Fig. 6, A and B). CFU granulocyte, erythroid, monocyte/macrophage, megakaryocyte (CFU-GEMM) progenitors did not display any differences between the two treatment groups, nor did the more mature lineage-restricted progenitors CFU-granulocyte/macrophage (CFU-GM) (Fig. EV6A).

**Figure 6.**
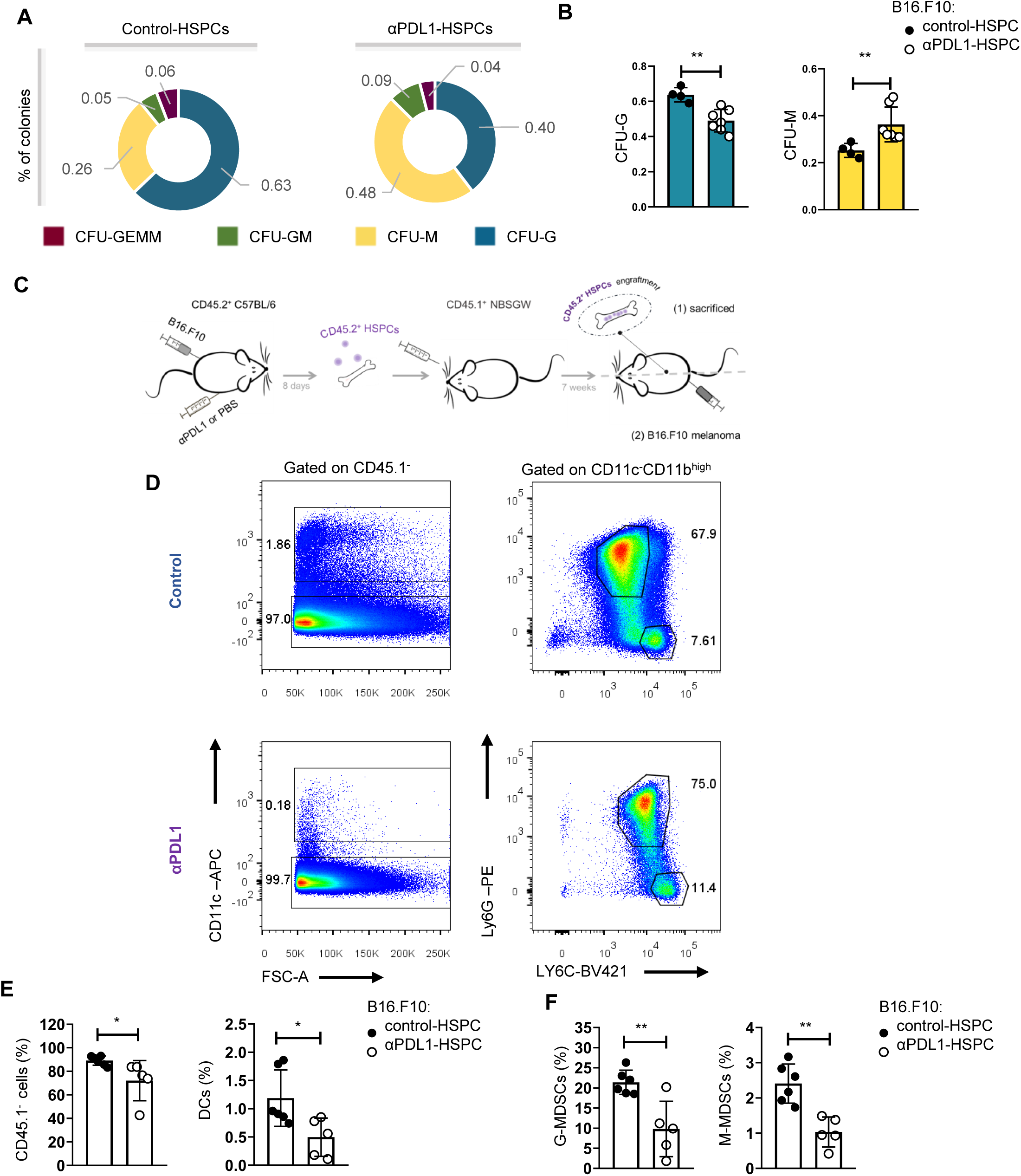
αPDL1 immunotherapy rewires cancer emergency myelopoiesis. (**A**-**B**) Representative CFU assay percentages (**A**) of cultured BM HSPCs isolated from B16.F10 inoculated C57BL/6 mice treated with PBS (control-HSPC) or αPDL1 (αPDL1-HSPC), sacrificed on the eighth day of tumour progression. Normalized frequencies of CFU-G, and CFU-M (**B**) of the control (n = 4) and αPDL1 (n = 7) conditions from 4 combined independent experiments. (**C**) Experimental scheme: HSPCs isolated from eight-days B16.F10 tumour-bearing C57BL/6 (CD45.2^+^CD45.1^-^) mice treated with PBS (control-HSPC) or αPDL1 (αPDL1-HSPC) and then adoptively transferred to NBSGW mice (CD45.2^-^CD45.1^+^). Following seven weeks of HSPC engraftment, recipient mice were either sacrificed and analyzed (Figure **C**; 1) or inoculated with B16.F10 and sacrificed after seventeen days of tumour development (Figure **C**; 2). (**D**; Numbers denote percentages of gated populations) Representative FACs plots of splenic DCs cells, G-MDSCs, and M-MDSCs isolated from melanoma bearing NBSGW (as in Figure 5 **C**). (**E**-**F**) Frequencies of splenic CD45.1^-^ cells, DCs (**E**; n = 6 control-HSPC, n = 5 αPDL1-HSPC), and in the CD11c^-^ population frequencies of CD11b^high^Ly6C^-^Ly6G^+^ G-MDSCs, and CD11b^high^Ly6C^+^Ly6G^-^ M-MDSCs (**F**; n = 6 control-HSPC, n = 5 αPDL1-HSPC) in NBSGW mice inoculated with B16.F10 and sacrificed after seventeen days (as in Figure 5C). Data from 2 combined independent experiments. p < 0.05*, p < 0.01**, p < 0.001***, p < 0.0001****. If not stated otherwise, unpaired two-tailed t-tests are performed. Means and SEM are depicted in all bar plots. n = biologically independent mouse samples

To provide direct evidence for an αPDL1-mediated altered myelopoiesis *in vivo,* we performed a transplantation experiment, as depicted in (Fig. 6C), where BM CD45.2^+^ HPSCs isolated from αPDL1- or PBS-treated, Β16.Β10 tumour-bearing mice were transplanted into CD45.1^+^ NOD.Cg-*Kit^W-41J^Tyr* ^+^*Prkdc^scid^ Il2rg^tm1Wjl^*/ThomJ (NBSGW) hosts, which support the multilineage engraftment of hematopoietic cells. The analysis of myeloid population composition of the spleen seven weeks after transplantations showed successful engraftment and comparable myelopoiesis potential between both groups (Fig. EV6B-D). Importantly, inoculation of mice with B16.F10 tumour cells demonstrated a rewiring of the myelopoiesis potential of HSPCs, derived from αPDL1-treated donor mice, as evident by the decreased frequencies of myeloid cells, including DCs, CD11b^high^ myeloid cells, and both MDSC subsets (Fig. 6D-F). Collectively, our findings demonstrate that αPDL1 immunotherapy alters the myelopoiesis program of HSPCs, altering their susceptibility to cancer-induced myelopoiesis.

## Discussion

Blockade of the PDL1/PD1 axis constitutes a highly promising therapy in a broad spectrum of solid tumours, which however, elicits durable anti-tumour responses and long-term remissions only in a small subset of patients (24, 25). Despite major research efforts, current biomarkers of response, such as the tumour mutational burden (TMB), the PD-L1 expression, the T cell infiltration, and IFNγ expression (26) demonstrate very low prediction power. For example, a recent meta-analysis showed that high TMB predicts responsiveness to αPDL1 only in 25% of patients with various types of cancer (27). Similarly, lack of PD-L1 expression cannot reliably exclude responses to αPDL1 or αPD1 ICΒ (28). Another major challenge is that clinical responses to PD1/PDL1 blockade are often accompanied by the development of adverse events resembling autoimmune reactions (29). Therefore, to understand the resistance mechanisms to immunotherapy and to design rational immunotherapies in cancer with diminished adverse events, it is necessary to delineate the unappreciated mechanisms of PD1/PDL1 axis targeted therapy. To this direction, herein we demonstrate that αPDL1 immunotherapy targets the HSPC compartment in the BM and rewires the cancer emergency myelopoiesis.

Since PD-1 engagement to PD-L1 imprints on T cell function to maintain tolerance (30), it was reasonable to focus on T cell-mediated anti-tumour immune responses as a potential mode of action of PD1/PDL1 targeting. However, recent studies demonstrate a broader expression of PD-1 and PD-L1, which adds a level of complexity to the so far proposed mechanisms. For example, PD-1 is also expressed by monocytic lineage cells, whereas PD-L1 is expressed by CD8 T cells, fibroblasts, endothelial cells (31–33). Our findings demonstrate that HSPCs express PD-L1 at steady state, and that it is upregulated depending on tumour immunogenicity. Although previous studies have shown the expression of PD-L1 by HSPCs (32, 34, 35), in this study we demonstrate that not only are they targeted by αPDL1 immunotherapy, but also that this interaction alters their fate and differentiation program. Specifically, αPDL1 immunotherapy promotes the expansion of HSPCs and induces their exit from quiescence, which is further supported by experiments with genetic ablation of the PD1/PDL1 axis. Molecularly we showed that inflammatory signalling is activated by treatment with αPDL1. Indeed, inflammatory signalling like IFN, S100A, TLR, IL-1, TNF is well established to promote activation and differentiation (35–40). Functionally this is translated by an altered cancer-associated emergency myelopoiesis as shown by the reduced myeloid cell frequency upon transplantation of αPDL1-treated HSPCs.

Tumour-associated myeloid cells constitute a heterogeneous population of cells that dictate the fate of tumour development. Tumour-associated macrophages (TAMs) and neutrophils (TANs), MDSCs, and DCs are the most abundant cell of myeloid origin in the tumour microenvironment (TME) (41), and mainly exert a tumour-promoting function. It is established that the majority of those cells originate from the BM through emergency myelopoiesis, which is directed by the nature of tumour cells (42). The unique characteristic of tumour-associated emergency myelopoiesis is the emergence of immature myeloid cells with intense immunosuppressive activities (42). Extensive research endeavours are focused on the reprogramming of cancer-associated emergency myelopoiesis to improve immunological performances against tumours. This has proven challenging due to myeloid cell heterogeneity and plasticity. Importantly, targeting strategies are focused on “terminally” differentiated myeloid cells, while efforts to interfere with myelopoiesis in the BM are limited. Interfering with cancer-associated myelopoiesis has been shown to be beneficial for host-promoting tumour regression in tumour-bearing mice. For example, transcriptomic and epigenetic rewiring of myelopoiesis induced by b-glucans resulted in the generation of granulocytes with anti-tumour activities, and this effect was transmissible by BM transplantation to naïve recipient mice (43). In line with this, our results showed that transplantation of αPDL1-treated HSPCs resulted in the reprogramming of tumour-induced myelopoiesis, as evidenced by the reduced potential of HSPCs from mice treated with aPDL1 to generate MDSCs in tumour-bearing recipient mice. Whether αPDL1 acts intrinsically on HSPCs or extrinsic mechanisms also participate to promote HSPC reprogramming remains to be investigated. Antibodies against PD-L1 have been shown to induce reverse signalling upon binding to tumour cells (44) but also to DCs (45) and macrophages (46). Accordingly, we show that αPDL1 treatment targets HSPCs in the BM, raising the possibility of a reverse signalling operation in the rewiring of emergency myelopoiesis. However, since systemic administration of αPDL1 is known to interfere with adaptive immune responses, the contribution of extrinsic mechanisms, such as the release of inflammatory cytokines and soluble factors, in HSPC reprogramming cannot be excluded. In contrast, genetic or pharmacological inhibition of PD-L1, showed to suppress the development of inflammatory macrophage in a yolk sac organoid culture (47). However, it is unlikely that this *in vitro* system provides all the necessary signals to mimic the *in vivo* development of macrophages.

From a mechanistic point of view, our data support a potential role of lymphocytes in the rewiring of tumour emergency myelopoiesis since αPDL1 treatment of tumour-bearing *RAG1^-/-^*animals failed to induce expansion of HSPCs in the BM. Recent evidence shows that Tregs in the BM are essential in regulating HSCs quiescence, while specific ablation of BM Treg cells leads to the expansion of HSCs and colony formation *in vitro* (48). Considering that Treg cells express high levels of PD-1, combined with our results showing that interruption of PDL1/PD1 axis leads to expansion of HSPCs in tumour-bearing mice, it is plausible that Treg cells may co-ordinate the cancer-associated emergency myelopoiesis through the PD-1 axis. Although the crosstalk of T cell subsets with HSCs has been previously reported (49, 50), whether this is supported by the PD1/PDL1 axis remains to be determined. Interestingly, another study showed that myeloid cell-specific ablation of PD-1 altered the emergency myelopoiesis, with myeloid progenitors such as CMPs and GMPs expressing high levels of PD-1 (32). Thus, the involvement of myeloid cells or stromal cells expressing PD-1 in shaping HSPC quiescence cannot be excluded. Of interest, in addition to PD-1, an interaction of PD-L1 with CD80 has been reported in mouse models (51). Both activated T cells (52) and myeloid cells express CD80 (53, 54); therefore, better characterization of the mechanisms that govern PDL1-mediated HSPC quiescence is required.

Treatment with αPDL1 caused transcriptomic reprogramming of HSPCs with the upregulation of inflammatory pathways. Indeed, these findings imply that the nature of myeloid cells that exit the BM upon αPDL1 treatment may possess an anti-tumour/inflammatory activity rather than a pro-tumourogenic/suppressive function. Accordingly, ablation of the PDL1/PD1 axis changed the balance of myeloid cells’ exit the BM with reduced MDSCs and increased effector myeloid cell frequencies (32). Thus, our findings may also hold important implications in the emergence of irAEs observed in patients responding to ICB immunotherapy. Although activation of effector CD4^+^ and CD8^+^ T cells (55, 56) as well as disturbances in Treg cells (57, 58) are implicated in irAEs development, inflammatory monocytes are expected to contribute in a direct way through the secretion of pro-inflammatory cytokines and chemokines or indirectly via antigen processing and presentation. However, the role of emergency myelopoiesis in irAEs has not been examined.

To conclude, our findings reveal the targeting of the HSPC compartment in the BM by αPDL1 immunotherapy, which reprograms the cancer-associated emergency myelopoiesis. Considering that PDL1/PD1 axis constitutes a major therapeutic target in solid tumours and hematologic malignancies, our data provide significant insights into therapy resistance mechanisms and the development of immune adverse events. Finally, the results described here place the BM microenvironment as a target of ICB immunotherapy for the future design of rational immunotherapy for cancer treatment.

## Materials and Methods

### Experimental Model and Subject Details

#### Human subjects

Seven patients with the diagnosis of Hodgkin’s disease were enrolled in this study; Five of them did not receive any treatment, and their BM samples were used for analysis of PD-L1 expression with flow cytometry. Two samples from the untreated patients were additionally used for RNAseq of isolated CD34^+^ cells together with samples from two patients with relapsed disease that received nivolumab. During BM collection, both patients under treatment were in remission.

#### Animals

C57BL/6J, and *Rag1^−/−^* mice (C57BL/6J background), NBSGW, and *PD-1^-/-^* mice were purchased from the Jackson Laboratory. The NBSGW humanized mouse strain was maintained as homozygotes (*NOD.Cg-Kit^W-41J^Tyr*^+^*Prkdc^scid^Il2rg^tm1Wjl^/ThomJ*). *Foxp3^EGFP^*.KI mice (C57BL/6 background) were kindly provided by A. Rudensky (Memorial Sloan–Kettering Cancer Center). Mice were housed 6 per cage in a temperature (21-23°C) and humidity-controlled colony room, maintained on a 12 hr light/dark cycle (07:00 to 19:00 light on), with standard food (4RF21, Mucedola Srl, Italy) and water provided *ad libitum* and environmental enrichments. All mice in the animal facility were screened regularly by using a health-monitoring program, in accordance to the Federation of European Laboratory Animal Science Association (FELASA) and were free of pathogens. All mice were maintained in the animal facility of the Biomedical Research Foundation of the Academy of Athens (BRFAA) and Institute of Molecular Biology and Biotechnology Institute (IMBB). During all experiments mice were monitored daily. All mice used in the experiments were female 8-12 weeks old. The total number of mice analyzed for each experiment is detailed in each figure legend. Littermates of the same genotype were randomly allocated to experimental groups.

#### Cell lines and primary cell culture

The B16.F10 melanoma and MB49 bladder cancer cell lines, used for the solid tumour induction models were kindly provided by A. Eliopoulos (Medical School, National and Kapodistrian University of Athens, Athens, Greece), and were negative for Mycoplasma spp., tested by PCR. B16.F10 and MB49 cancer cells were cultured at 37°C under 5% CO2 in RPMI-1640 (GlutaMAX™, Gibco, #61870) and DMEM (Gibco, #11965) medium, respectively, supplemented with 10% heat-inactivated fetal bovine serum (FBS, Gibco, #10270), 100 U/ml penicillin–streptomycin (10,000 U/ml, Gibco, #15140), and 50 μM 2-mercaptoethanol (50 mM, Gibco, #31350). Cells were split at 90%–100% confluence. All experiments were performed with early passage (p2–3) cells.

Murine sorted MDSCs and Teff cells were obtained as described below. They were cultured in RPMI-1640 medium containing 10% heat-inactivated FBS, 100 U/ml penicillin–streptomycin, and 50 μM 2-mercaptoethanol.

#### Solid tumour induction and in *vivo* immunotherapy administration protocols

Mice were implanted subcutaneously (s.c.) on the back with 3 × 10^5^ B16.F10 melanoma (59) or 75 × 10^4^ MB49 bladder cancer cells. Cancer cells viability was assessed by Trypan blue exclusion. Mice were then euthanized on indicated in the date each experimental setting. Mice that manifested tumour ulceration were excluded for the experimental processes.

For the application of immunotherapy, mice were treated intraperitoneally (i.p.) with anti-PD-L1 (αPDL1) antibody (αPDL1; 200 μg per 100 μl; i.p: clone MIH5), anti-CTLA-4 (100 μg per 100μl in each mouse i.p. : clone 4F10), anti-PD-1 (αPD1; 200 μg per 100 μl i.p. : clone RMP1–14). Control mouse cohort was administered i.p. PBS on the same days. Immunotherapy or control treatment were administered every 3 days after tumour implantation.

#### Tissue dissociation and sample preparation

Lymph nodes and spleen were collected from euthanized mice, and single-cell suspensions were obtained by homogenization of the tissues and filtering through a 40-μm cell strainer (BD Falcon) with ice cold 5% FBS/PBS. Tibiae, femurs, and hip bones were collected, and BM cell suspension was isolated by flushing out the bones with ice cold 5% FBS/PBS. Red blood cells in spleen and BM cell suspensions were lysed by incubation in 2 ml Ammonium chloride (NH_4_Cl) for 2 min in RT. Cells from the TME were isolated by dissociating tumour tissue in the presence of RPMI-1640 (GlutaMAX™, Gibco, #61870) supplemented with collagenase D (1 mg ml^−1^, Roche) and DNase I (0.25 mg ml^−1^, Sigma) for 45min before passing through a 40-μm cell strainer (BD Falcon). Peripheral blood collection was obtained through the collection the submandibular vein using 25-gauge needle. To prevent blood from clotting, a solution of 0.1 M EDTA was used for coating syringes, needles and tubes. PBMCs were isolated on Lymphocyte Separation Media (Lymphosep; biowest, #L0560). Tubes were centrifuged at 500 g for 30 min with no brake RT. PBMC layer was collected, and cells were washed with PBS.

Human BM aspirates were collected from patients from the HL patients and BM mononuclear cells were isolated by density gradient centrifugation, using Ficoll-Histopaque 1077 (Sigma-Aldrich, #10771).

#### Flow cytometry and cell sorting

For extracellular marker staining, single-cell suspensions from murine tumour, spleen, LNs, peripheral blood or BM were incubated for 20 min at 4°C with the following anti-mouse conjugated antibodies: anti-CD45-PerCP/Cy5.5 (BioLegend, clone 30-F11, #103132), anti-CD45.1-PE/Cyanine7 (BD Biosciences, clone A20, #560578), anti-CD11c-APC (BioLegend, clone N418, #117310), anti-CD11c-FITC (BioLegend, clone N418, #117306), anti-CD11c-PE (BioLegend, clone N419, #117308), anti-CD11b-PE/Cyanine7 (BioLegend, clone M1/70, #101216), anti-CD11b-BrilliantViolet 510 (BioLegend, clone M1/70, #101263), anti-CD11b-FITC (BioLegend, clone M1/70, #101206), anti-Gr-1-PE (BioLegend, clone RB6-8C5, #108408), anti-Gr-1-PE/Cyanine7 (BioLegend, clone RB6-8C5, # 108416), anti-Gr-1-FITC (BioLegend, clone RB6-8C5, #108406), anti-Ly-6G-PE (BioLegend, clone 1A8, #127608), anti-Ly-6G-PE/Cyanine7 (BioLegend, clone 1A8, # 127618), anti-Ly-6C-BrilliantViolet421 (BioLegend, clone HK1.4, #128032), anti-Ly-6C-PerCP (BioLegend, clone HK1.4, # 128028), anti-CD274 (B7-H1, PD-L1)-Brilliant Violet 421 (BioLegend, clone 10F.9G2, #124315), anti-CD274 (B7-H1, PD-L1)-Brilliant Violet 421 (BD Pharmingen, clone MIH5, # 564716), anti-CD274 (B7-H1, PD-L1)-PE/Dazzle (BioLegend, clone, 10F.9G2, #124323), anti-TER-119-FITC (BioLegend, clone TER-119, #116206), anti-CD45R/B220-FITC (BioLegend, clone RA3-6B2, #103206), anti-CD16/32-FITC (BioLegend, clone 93, #101306), anti-CD16/32-PE/Cyanine7( BioLegend, clone 93, # 101317), anti-CD117 (c-Kit)-PE (BioLegend, clone 2B8, #105808), anti-Ly-6A/E (Sca-1)-APC (BioLegend, clone E13-161.7, #122512), anti-Ly-6A/E (Sca-1)-Brilliant Violet 421 (BioLegend, clone D7, #108127), anti-CD48-Alexa Fluor 700 (BioLegend, clone HM48-1, #103426), anti-CD150-PE/Cyanine7 (BioLegend, clone TC15-12F12.2, #115914), anti-CD34-Brilliant Violet 421 (BioLegend, MEC14.7, #119321), anti-CD135-Brilliant Violet 421 (BioLegend, clone A2F10, #135313), anti-CD127 (IL-7Rα)-PerCP/Cyanine5.5 (BioLegend, clone SB/199, #121114), anti-CD3ε-Pacific Blue (BioLegend, clone 145-2C11, #100334), anti-CD4-Brilliant Violet 510 (BioLegend, clone GK1.5, #100449). FMO was used as a negative control, to increase the accuracy of gate placement. Data acquisition was performed on a FACSAria III (BD Biosciences), FACSCelesta (BD Biosciences), FACS Canto II (BD Biosciences) and the BD FACSDiva v8.0.1 software (BD Biosciences). Murine splenic MDSCs, and T effector cells, and BM HSPCs were sorted on a FACS ARIA III (BD Biosciences) v8.0.1 software (BD Biosciences). Cell purity was above 95%. Flow cytometry data were analyzed with FlowJo v.8.7 and 10.8.1 software.

Human BM mononuclear cells were stained for extracellular surface markers, in staining buffer (2% FBS/PBS) for 20 min at 4°C, before acquisition via flow cytometry. The following human monoclonal antibodies were used: anti-CD34-FITC (BioLegend, clone 581, # 343504), anti-CD34-APC (BD Biosciences, clone 8G12, # 345804), anti-PD-L1-PE (BioLegend, clone 29E.2A3, # 329705), anti-CD45-PerCP (BD Biosciences, clone 2D1, # 347464), anti-CD38-APC-H7 (BD Biosciences, clone HB7, # 653314). FMO was used as a negative control, to increase the accuracy of gate placement. Cell acquisition was performed with a FACS Canto II flow cytometer (BD Biosciences) and cells were sorted on a FACS ARIA III (BD Biosciences) v8.0.1 software (BD Biosciences). Cell purity was above 95%.

#### Cell cycle assessment

For the cell cycle analysis via flow cytometry 10^6^ BM HSPCs per sample were first stained extracellularly, as previously described, fixed and permeabilized using fixation / permeabilization buffer (Foxp3/TF Buffer Set; eBioscience, #00552300) and subsequently stained with Ki-67-PE/Cyanine7 (BioLegend, clone 16A8, # 652425). At the end, cells were stained with 7-AAD (BioLegend, #420404) cellular DNA content marker (5 μl per sample) in 200 μl of 5% FBS/PBS. Cells were analyzed using BD FACSCelesta using the BD FACSDiva v8.0.1 software. Linear scale was used for 7-AAD.

### HSPC transplantation

CD45.2^+^-C57BL/6 mice were inoculated with B16.F10 melanoma cells and were treated with αPDL1 or PBS as previously described. Eight days after injection, mice were euthanized, BM cells were isolated as previously described, and 2 x 10^4^ HSPCs were injected in the orbital vein of humanized mice CD45.1^+^-NBSGW. 6 -7 weeks post injection, NBSGW mice were either sacrificed or subcutaneously injected with B16.F10, as previously described. After 17 days, NBSGW mice were euthanized, and the lineage output was measured through flow cytometry.

### Colony Forming Unit Assay (CFU)

4.000 sorted BM HSPCs from B16.F10 inoculated mice, control or αPDL1 treated, were seeded in complete Methocult (Stem Cell Technologies, M3434, # 03434), which was plated in 35-mm tissue culture dishes (Sigma, CLS430165). The enumeration and identification of colonies (CFU-M, CFU-G, CFU-GM, CFUGEMM) was scored after 8-12 days in culture (at 37°C in 5% CO_2_) under an inverted light microscope according to the manufacturer’s instructions. For every biological mouse sample, technical duplicate cultures were seeded. Each type of colony is represented normalized, derived from the equation:

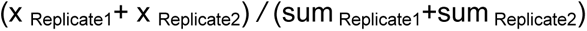

where “x” is the sum identified for each colony per replicate and “sum” the total number of colonies for each replicate, regardless of identity.

### *In vitro* suppression assay

For the suppression assay of MDSC subsets, CD4^+^Foxp3^-^ effector T cells (Teff) were sorted from the LNs of naïve Foxp3^EGFP^.KI mice, as described previously (60, 61), and stained with the division-tracking dye CellTrace CFSE (Invitrogen, #C34554) according to the manufacturer’s protocol. In summary, a total of 75 × 10^3^ labeled Teff cells were seeded in 96-well round-bottom plate in each well supplemented with Dynabeads™ Mouse T-Activator CD3/CD28 (Gibco, #11456D) at ration 1:1 beads to Teff cells. M-MDSC (CD11b^high^Ly6C^+^Ly6G^−^) and G-MDSC (CD11b^high^Ly6C^−^Ly6G^+^) subsets, sorted from the spleens of C57BL/6J B16.F10 inoculated mice treated with either αPDL1 or control (PBS), were added at the culture for a total of 64 hr at a ratio Teff/M-MDSCs 1:1 and Teff/G-MDSCs 3:1.

### BM fluid isolation and preparation for proteomic analysis

Femurs were isolated and were flushed with ice cold PBS in Eppendorf tubes. The BM supernatant was harvested after pelleting cells by centrifugation at 1800 rpm for 10 min at 4°C. Cytokine profile was evaluated via Mass Spectrometry.

Bone marrow supernatants (0.5 mL per sample) were concentrated with 3 kDa MWCO Amicon Ultra Centrifugal filter devices (Merck Millipore) up to a final volume of 30 μL. Protease inhibitors were added to the samples and the protein concentration was defined with Bradford Assay. Concentrated samples were processed with the filter aided sample preparation (FASP) method as described previously (62), with minor modifications (63). Briefly, sample volume corresponding to 200 μg of total protein content was mixed with lysis buffer (0.1 M Tris-HCl pH 7.6, supplemented with 4% SDS and 0.1 M DTE) and buffer exchange was performed in Amicon Ultra Centrifugal filter devices (0.5 mL, 30 kDa MWCO; Merck Millipore) at 14,000 rcf for 15 min at RT. Each sample was diluted with urea buffer (8 M urea in 0.1 M Tris-HCl pH 8.5) and centrifuged. The concentrate was diluted again with urea buffer and centrifugation was repeated. Alkylation of proteins was performed with 0.05 M iodoacetamide in urea buffer for 20 min in the dark, RT, followed by a centrifugation at 14,000 rcf for 10 min at RT. Additional series of washes were conducted with urea buffer (2 times) and ammonium bicarbonate buffer (50 mM NH_4_ HCO_3_ pH 8.5, 2 times). Tryptic digestion was performed overnight at RT in the dark, using a trypsin to protein ratio of 1:100. Peptides were eluted by centrifugation at 14,000 rcf for 10 min, lyophilized and stored at –80°C until further use.

### LC-MS/MS analysis

Samples were resuspended in 200 μL mobile phase A (0.1% FA). A 5 μL volume was injected into a Dionex Ultimate 3000 RSLS nano flow system (Dionex, Camberly, UK) configured with a Dionex 0.1 × 20 mm, 5 μm, 100 Å C18 nano trap column with a flow rate of 5 µL / min. The analytical column was an Acclaim PepMap C18 nano column 75 μm × 50 cm, 2 μm 100 Å with a flow rate of 300 nL / min. The trap and analytical column were maintained at 35°C. Mobile phase B was 100% ACN:0.1% Formic acid. The column was washed and re-equilibrated prior to each sample injection. The eluent was ionized using a Proxeon nano spray ESI source operating in positive ion mode. For mass spectrometry analysis, a Q Exactive Orbitrap (Thermo Finnigan, Bremen, Germany) was operated in MS/MS mode. The peptides were eluted under a 120 min gradient from 2% (B) to 80% (B). Gaseous phase transition of the separated peptides was achieved with positive ion electrospray ionization applying a voltage of 2.5 kV. For every MS survey scan, the top 10 most abundant multiply charged precursor ions between m/z ratio 300 and 2200 and intensity threshold 500 counts were selected with FT mass resolution of 70,000 and subjected to HCD fragmentation. Tandem mass spectra were acquired with FT resolution of 35,000. Normalized collision energy was set to 33 and already targeted precursors were dynamically excluded for further isolation and activation for 15 sec with 5 ppm mass tolerance.

### MS data processing

Raw files were analyzed with Proteome Discoverer 1.4 software package (Thermo Finnigan), using the Sequest search engine and the Uniprot mouse (Mus musculus) reviewed database, downloaded on November 22, 2017, including 16,935 entries. The search was performed using carbamidomethylation of cysteine as static and oxidation of methionine as dynamic modifications. Two missed cleavage sites, a precursor mass tolerance of 10 ppm and fragment mass tolerance of 0.05 Da were allowed. False discovery rate (FDR) validation was based on q value: target FDR (strict): 0.01, target FDR (relaxed): 0.05.

Normalized serum protein concentrations were imported into R, sample mean fluorescence intensities were scaled to each other, log2 transformed, and plotted in a heatmap using the heatmap.2 function from the gplots package package v3.1.1. Proteins were considered differentially abundant at a cutoff of |FC| ≥ 1.5, and significant at P < 0.05, as determined by unpaired two-tailed Student’s t-test. Functional enrichment analysis tables of significant differentially abundant proteins were produced with Metascape v3.5 (http://metascape.org), and top hits were visualized in a dot-plot using the R ggplot2 package v3.4.1. Gene set enrichment analysis (GSEA v4.2.2 [build: 8]) was performed to reveal enriched signatures in our gene sets based on the Molecular Signatures Database (MSigDB, v7.4). Gene sets were ranked by taking the –log10 transform of the p-value and multiplying it by the corresponding FC, with significantly upregulated genes at the top of the ranked list. GSEA pre-ranked analysis was performed using the remapped Mouse Gene Symbol dataset and collapsing probe sets, while keeping only the max probe value. The rest of the parameters were left to default. Enrichment was considered significant if FDR (q-value) < 25%. Pathway analysis was performed using tissue specific Ingenuity Pathway Analysis (IPA Winter Release (Dec 2022), RRID:SCR_008653).

### RNA sequencing library preparation

BM murine samples were isolated from B16.F10 melanoma, and MB49 bearing mice treated with PBS or αPDL1. Human BM samples, as previously described, were from HL patients isolated at diagnosis or after αPD1 treated. HSPCs were sorted, and total RNA was extracted using Arcturus™ PicoPure™ RNA Isolation Kit (Thermo Fisher Scientific, #12204-01) ’ instructions.

B16.F10-bearing and MB49 HSPCs and Human BM-derived CD34^+^ cells RNAseq experiments were carried out at the Greek Genome Center (GGC) of the Biomedical Research Foundation of the Academy of Athens (BRFAA). RNAseq libraries were prepared with the NEBNext Ultra II Directional RNA Library Prep Kit for Illumina (Illlumina), Quality Control was performed with the Agilent bioanalyzer DNA1000 kit and Quantitation with the qubit HS spectrophotometric method. Approximately 25 Million 100 bp single-end reads were generated for each sample in the Illumina Novaseq 6000 system.

MB49 bladder cancer RNA-seq library preparation was carried out at the Max Planck Institute of Immunobiology and Epigenetics (MPI-IE). cDNA libraries were prepared using SMART-seq® v4 Ultra Low Input RNA Kit (#634888, Takara). The NEB Ultra II FS DNA kit (#E7805S) was used to generate barcoded sequencing libraries. Quality control was performed with Agilent 5200 Fragment Analyzer. 50 million paired-end 101 bp reads per sample were generated using the Illumina Hiseq 3000 or NovaSeq 6000 system, at the DeepSequencing Facility at MPI-IE

### RNA sequencing data processing

RNA-seq data analysis paired-end fastq read files were pre-processed by assessing for quality with FastQC v0.11.9 and trimming off Illumina sequencing adapters with galore Trim Galore v0.3.7. Alignment to the reference mouse genome (GENCODE GRCm38.p6_M230) was carried out with STAR v2.7.10a using the default parameters. HTseq-count v0.12.4 was used to produce gene count matrices from the resulting alignments with the specific parameters -- *intersection-non-empty*, and additionally *--stranded=”reverse”* (using the reference GENCODE GRCm38.p6_M23 annotation) for RNA libraries prepared with the NEB directional kit. Sample hierarchical clustering and PCA, TMM normalization, scaling, and differential expression analysis via exact test was performed in R using edgeR v3.34.1. Mouse genes were considered significantly differentially expressed if they met |FC| ≥ 1.5 and FDR< 0.05. Human DEGs were considered significant at FDR<0.05.

Functional enrichment analysis tables of DEGs were produced with g:Profiler version *e109_eg56_p17_1d3191d* web-server, and top hits were visualized as dot-plots using the R ggplot2 package v3.4.1. GSEA pre-ranked analysis and IPA was performed on the murine data sets, as described in proteomics.

### Data analysis and statistics

Data are presented as mean ± S.D., as bar graphs represent the mean and standard deviation (SD) between biologically independent mouse samples or technical replicates, as indicated in corresponding the figure legend. For statistical analysis, all data were analyzed using Prism 8 (GraphPad Software, Inc., La Jolla, USA). Data were analyzed using the two-tailed, parametric, unpaired Student’s t test or the two-tailed, nonparametric Mann–Whitney test, as appropriate after testing for normality of the values with the F test, with 95% confidence intervals. For multiple-group comparisons, the two-way ANOVA Tukey’s multiple comparison test was performed. The p-value < 0.05 was considered to be statistically significant for each dataset.

### Study approval

The study was approved by the institutional review board and ethics committee of G. Papanicolaou Hospital (135/2020). All patients gave written informed consent. The study was conducted in compliance with the Helsinki Declaration.

All mice were maintained in the animal facility of the BRFAA and IMBB. All procedures were in accordance with institutional guidelines and were approved by the Institutional Committee of Protocol Evaluation of the BRFAA and the Institutional Committee of Protocol Evaluation of the IMBB together with the Directorate of Agriculture and Veterinary Policy, Region of Attika, Greece (Athens, Greece 299868, 7/4/2022, and 557279, 30/07/2020) and the Directorates of Agricultural Economy and Veterinary, Region of Crete, Greece (Heraklion, Greece, 216160, 20/07/2022).

## Data availability

Human and mouse data are currently under submission. Human RNA-seq data in European Genome-Phenome Archive EGA ( https://ega-archive.org/). Mouse RNA-seq in GEO (https://www.ncbi.nlm.nih.gov/geo/) and mass spectrometry proteomic data in the ProteomeXchange Consortium via the PRIDE (https://www.proteomexchange.org/).

## Acknowledgements

We wish to thank Anastasia Apostolidou for her technical assistance on flow cytometry and cell sorting, Athina Varveri, Themis Alissafi, and Miranta Papadopoulou for assisting with experiments, Pavlos Alexakos for assisting with mice, Giannis Vatselas for assisting with RNA-seq and for providing technical advice. Support was provided by the German Research Foundation, grant nos. 322977937/GRK2344 (to E.T. and P.B.) and GZ TR 1478/2-1 (to E.T.), the grant no. FRM AJE202010012488 (to E.T.), the Labex Chair of excellence (to E.T.). E.G., I.M. and P.V. were supported by the General Secretariat for Research and Technology Management and Implementation Authority for Research, Technological Development and Innovation Actions (MIA-RTDI) (grant Τ2EDK-02288, MDS-TARGET).

## Author Contributions

A.B. performed experiments, analyzed data, generated figures, and wrote the manuscript, A.S.P and I.P. performed experiments and analyzed data, P.B., M.G and V.B. assisted with the transcriptomic and bioinformatic analysis, L.B. provided materials and assisted in designing experiments, M.M and A.V. performed the proteomic analysis, A.T. performed experiments with clinical samples, M.I., E.G. performed clinical evaluation of HL individuals, A.H. assisted with experiments and critically edited the manuscript, I.M. assisted with the cohort development, analyzed patient data and critically edited the manuscript, E.T. designed and supervised the transcriptomic analysis, provided critical insights in the design of the study P.V. designed and supervised the study, performed data analysis, and wrote the manuscript.

## Expanded View Figure legends

**Expanded View Figure 1.**
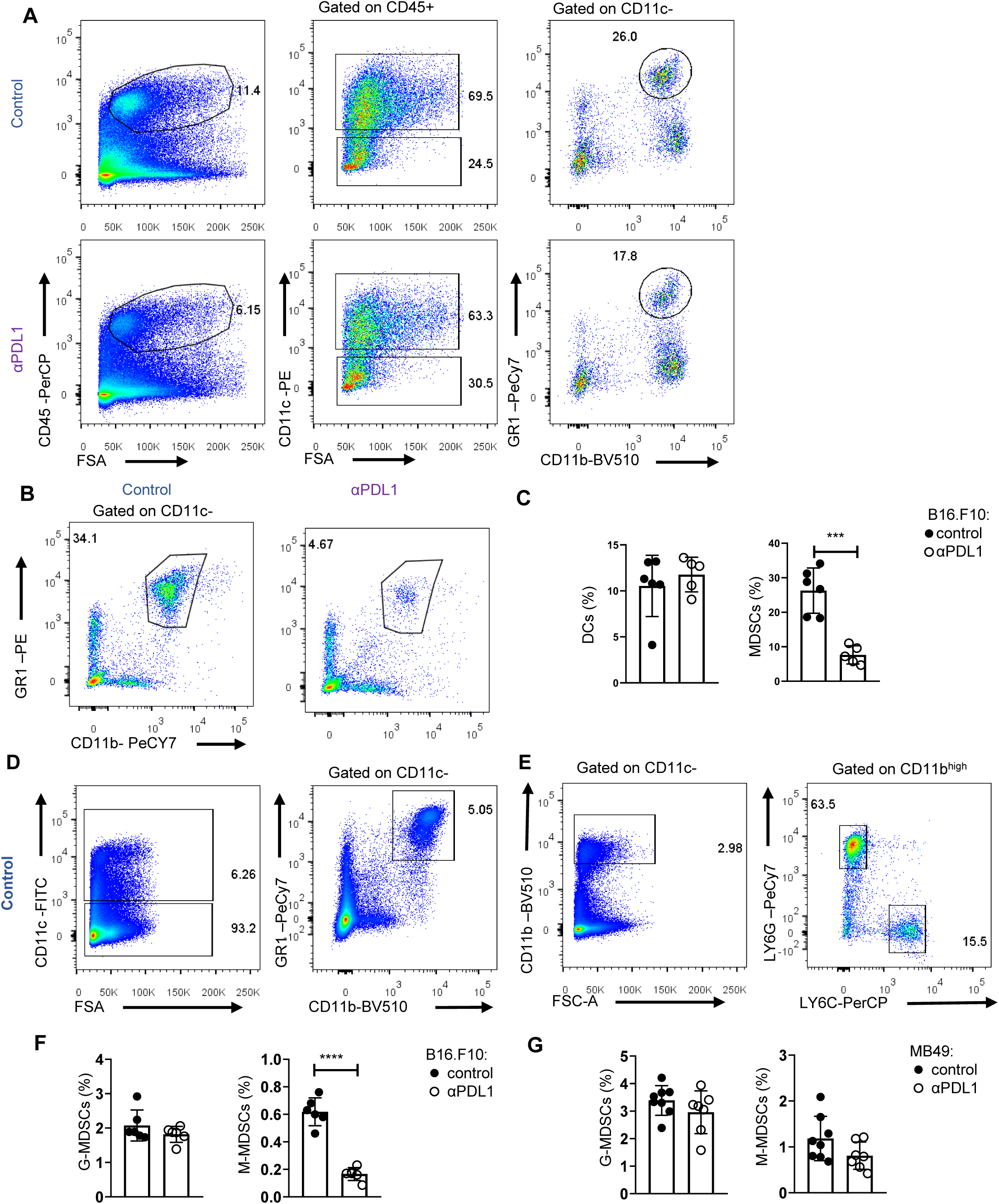
PD-L1 blockade results in a significant peripheral and intratumoural population reformatting during tumour progression. (**A**) Gating strategy of intratumoural CD45^+^ cells, DCs, MDSCs of PBS (control) or αPDL1 treated C57BL/6 mice sacrificed after eight days of B16.F10 tumour progression. Numbers denote percentages of gated populations. (**B**-**C**) Representative facs plots (**B**; Numbers denote percentages of gated populations) and quantification (**C**; n = 6 control, n = 5 αPDL1) of peripheral blood DCs, and MDSCs from eight days B16.F10 inoculated PBS or αPDL1 treated C57BL/6 mice. Representative data from 3 independent experiments, assessed with Flow cytometry. (**D**-**E**) Gating strategy of splenic DCs, MDSCs (**D**), M-MDSCs, and G-MDSCs(**E**) after eight days of B16.F10 inoculation of PBS or αPDL1 treated C57BL/6 mice. Numbers denote percentages of gated populations. (**F**-**G**) Frequencies assed in the CD11c^-^ population of splenic G-MDSCs, and M-MDSCs after eight days of B16.F10 (**F**; n = 6 control, n = 6 αPDL1), and MB49 (**G**; n = 8 control, n = 7 αPDL1) inoculation in C57BL/6 mice treated with PBS (control) or αPDL1. Representative data assessed with Flow cytometry from 2 independent experiments (**F**), and data from 2 combined independent experiments (**G**). p < 0.05*, p < 0.01**, p < 0.001***, p < 0.0001****. If not stated otherwise, unpaired two-tailed t-tests are performed. Means and SEM are depicted in all bar plots. n = biologically independent mouse samples.

**Expanded View Figure 2.**
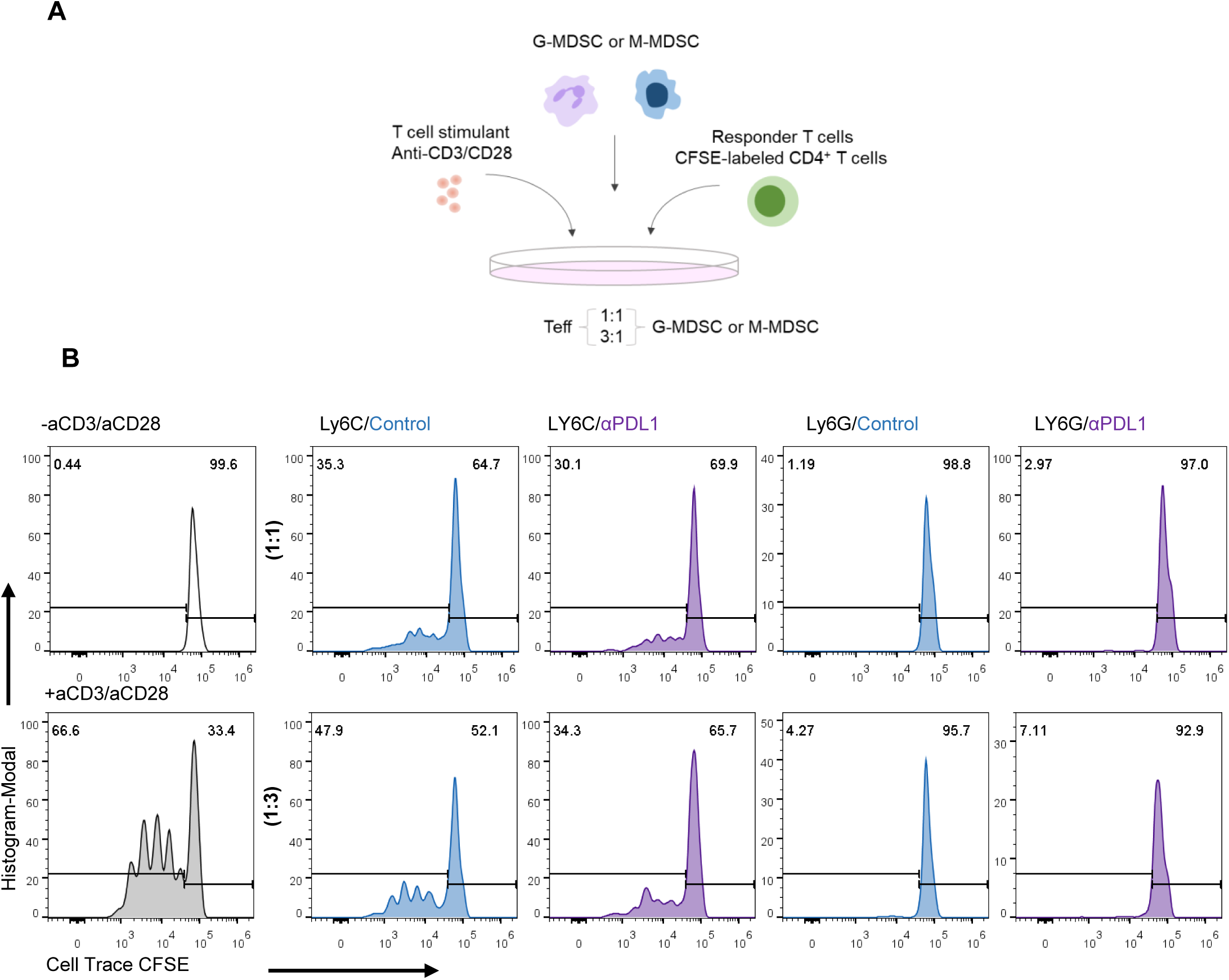
MDSC subsets from αPDL1 treated mice exhibit sustained suppressive function compared to control. (**A**) Experimental outline for the evaluation of the suppressive ability of MDSC subsets *in vitro*. G-MDSCs and M-MDSCs were sorted from the spleen of C57BL/6 B16.F10 bearing mice treated with PBS (n = 2) or αPDL1(n = 2) and sacrificed after eight days. Cells were co-cultured, at different ratios, activated by anti-CD3/anti-CD28 monoclonal antibodies Cell Trace Violet (CTV)-labeled CD4^+^Foxp3^−^ T effector cells, isolated from the spleens of Foxp3^EGFP^ naïve mice. Suppressive activity of MDSC subsets was estimated with flow cytometry after 62 hr. (**B**) Representative Histograms of CTV MFI dilution of different culture conditions (MDSC-subset: Teff 1:1, and 1:3). Teff cell proliferation is represented through the gating of histograms in CD4^+^CTV^+^ cells^+^. Percentages of proliferated cell numbers are represented in the numbers in the histograms. Representative data from 2 independent experiments (n = 2 mice/group in each experiment)

**Expanded View Figure 3.**
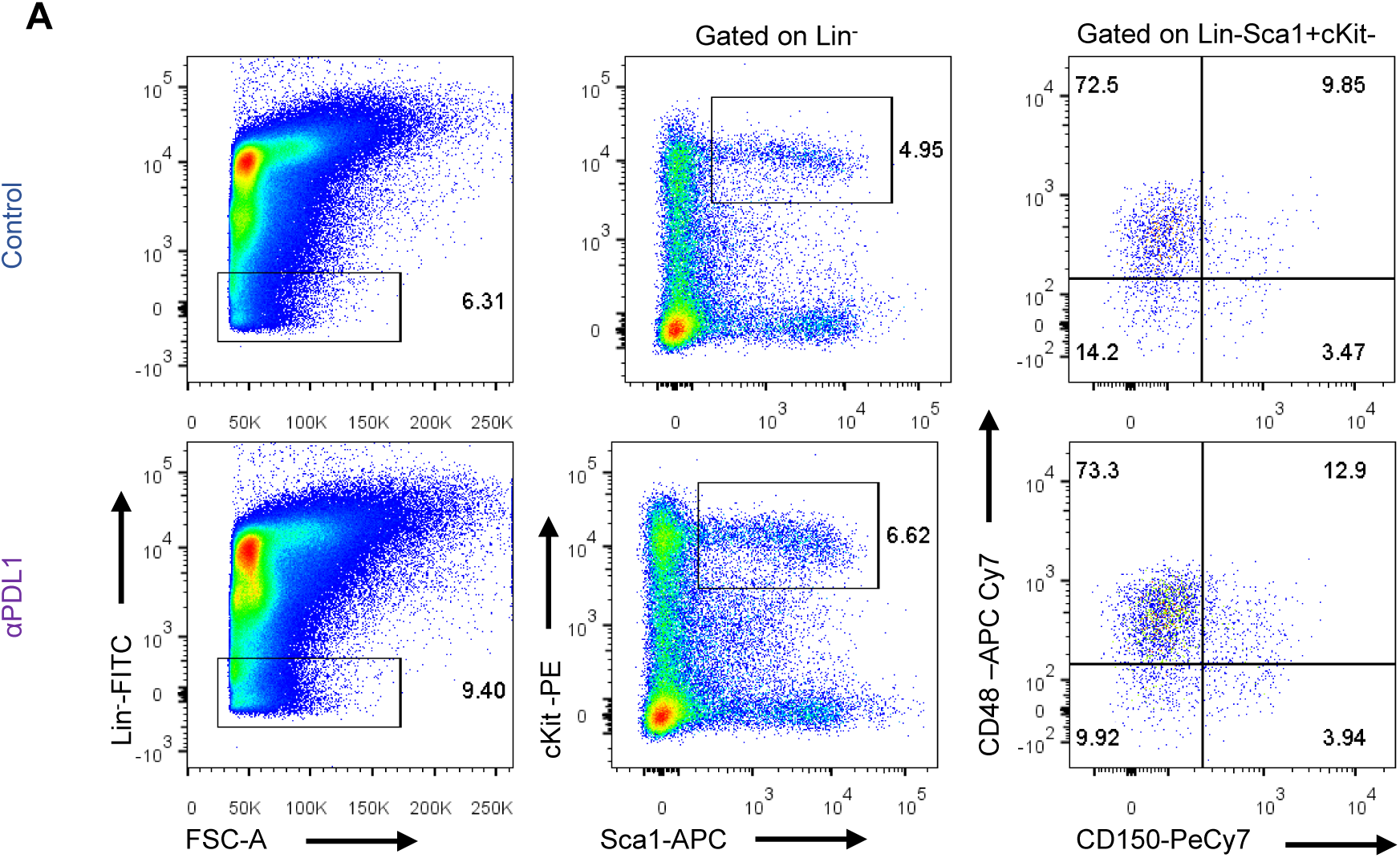
Gating strategy of BM HSPCs of control or αPDL1 melanoma bearing mice. (**A**) Gating strategy of BM HSPCs (LSK: Lin^-^Sca1^+^cKit^+^), including the three subpopulations LT-HSCs, ST-HSCs, and MPPs isolated from eight days B16.F10 tumour bearing C57BL/6 treated with PBS or αPDL1. Numbers denote percentages of gated populations

**Expanded View Figure 4.**
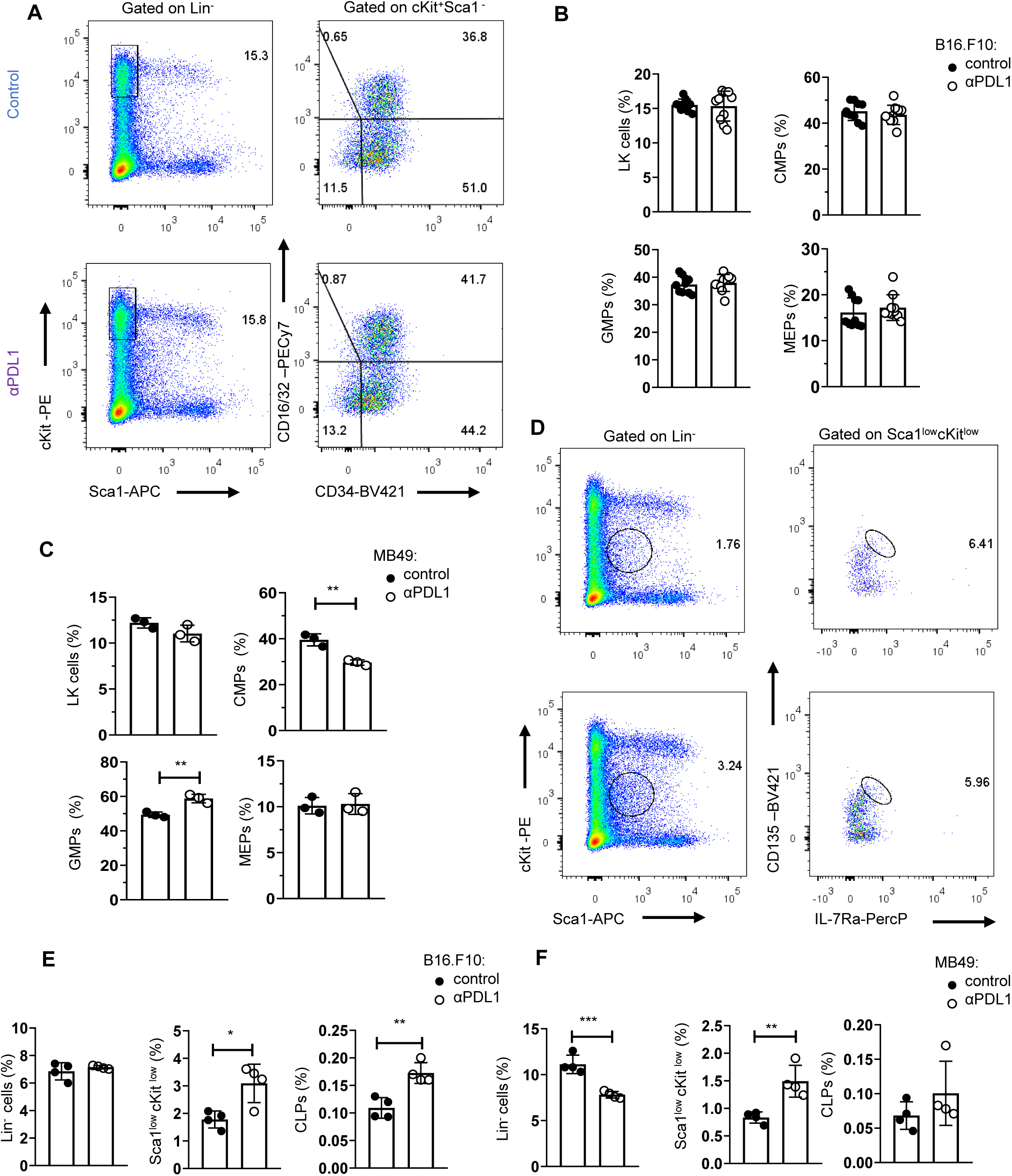
Tumour immunogenicity dictates the differentiation potential of αPDL1 targeted HSPCs. (**A**) Gating strategy of the BM LK pool (Lin^-^Sca1^-^cKit^+^), and the subclusters CMPs (LK CD34^+^CD16/32^-^), GMPs (LK D34^+^CD16/32^+^), and MEP (LK CD34^-^ CD16/32^-^) isolated from treated PBS or αPDL1 C57BL/6 mice inoculated with B16.F10 and sacrificed after eight days. Numbers denote percentages of gated populations. (**B**-**C**) Frequencies of the BM LK compartment, CMPs, GMPs, and MEPs isolated from treated PBS or αPDL1 C57BL/6 mice inoculated with melanoma (**B**; control n = 10, n = 10 αPDL1) or MB49 (**C;** n = 3 control, n = 3 αPDL1) and sacrificed after eight days. Representative data from 2 (**C**) and 2 combined out of 4 (**B**) independent experiments. (**D**-**F**) Gating strategy common lymphoid progenitors (CLPs) in the BM (Lin^-^Sca1^low^cKit^low^ IL-7Rα^hi^CD135^hi^) (**D**; Numbers denote percentages) and their frequencies in C57BL/6 mice treated with PBS or αPDL1 and inoculated with either B16.F10 (**E**; n = 4 control, n = 4 αPDL1) or MB49 (**F**; n = 4 n = 4 control, n = 4 αPDL1), sacrificed on the eight day of tumour development. of gated populations Representative data from 1 experiment (**F**) and 2 (**E**) independent experiments. p < 0.05*, p < 0.01**, p < 0.001***, p < 0.0001****. Means and SEM are depicted in all bar plots. If not stated otherwise, unpaired two-tailed t-tests are performed. n = biologically independent mouse samples

**Expanded View Figure 5.**
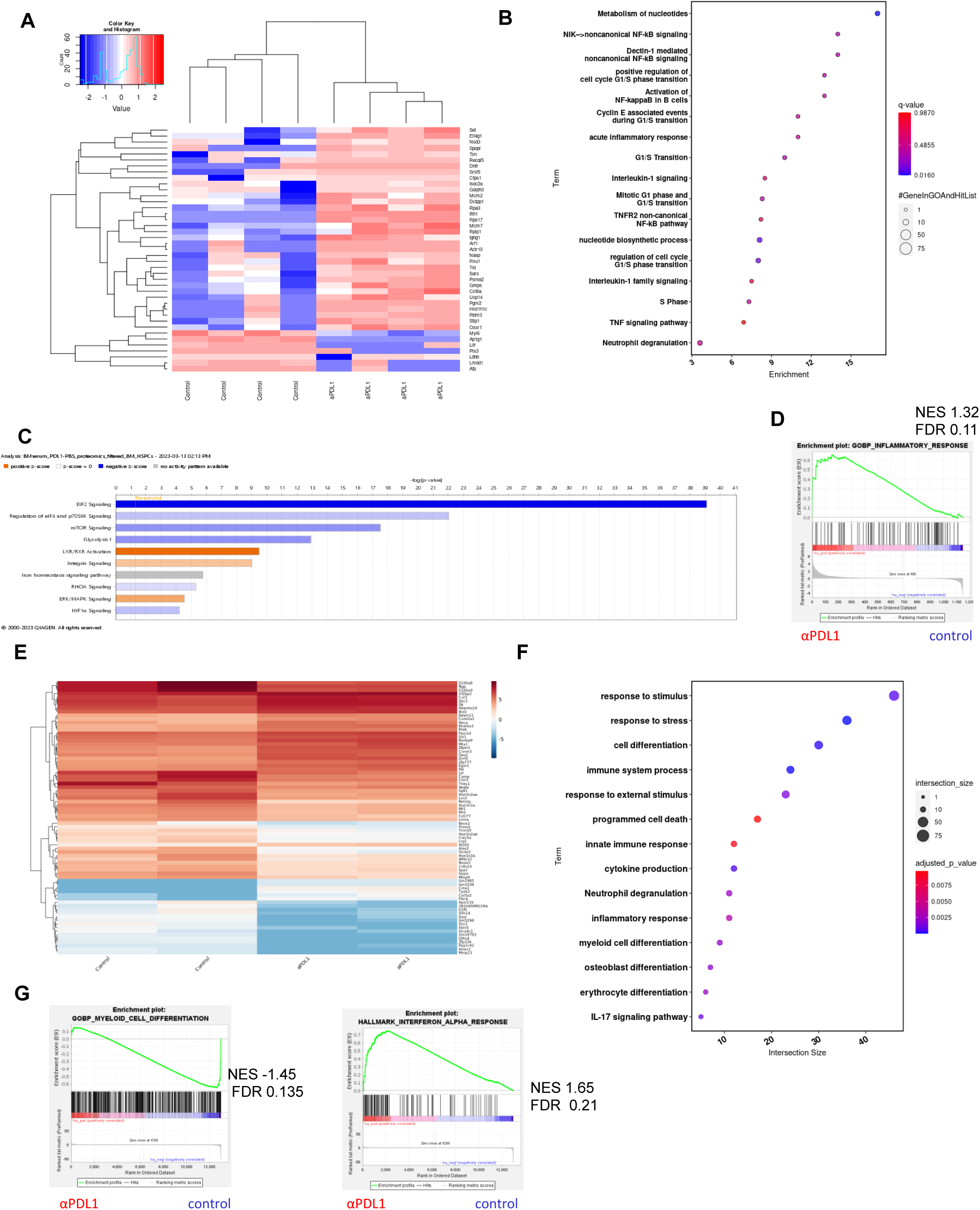
Proteomic and transcriptomic analysis of HSPCs corroborate towards a transcriptomic re-programming upon treatment with αPDL1 blockade. (**A**-**C**) BM aspirates from αPDL1 (n = 4) vs. PBS (control n = 4) treated B16.10 bearing C57BL/6 mice, sacrificed on the eight day of tumor development and used for proteomic analyses: (**A**) Heatmap of the 41 DEPs (7 up-regulated, and 34 down-regulated in αPDL1 vs control) which displayed a differential abundance (|FC| ≥ 1.5, p < 0.05). (**B**) Enrichment pathway analysis of significantly differentially abundant proteins. (**C**) BM specific Ingenuity pathway analysis (IPA). (**D**) GSEA plot showing the positive enrichment pathway “Inflammatory response” (NES 1.32, FDR 0.11) in αPDL1 compared to control. n = biologically independent mouse samples. (**E**-**G**) RNA-sequencing BM HSPCs isolated from MB49 tumour-bearing C57BL/6 mice treated with PBS (n = 2) or αPDL1 (n = 2) (**E**) Heatmap portraying the 76 DEGs (29 genes up-regulated, and 47 genes down-regulated in αPDL1 vs controls, |FC|>1.5, FDR<0.05). (**F**) Pathway analysis of the DEGs. (**G**) GSEA plot showing the negative enrichment of “Myeloid Cell Differentiation” (NES -1.45, FDR 0.135), and positive enrichment “Interferon alpha response” (NES 1.65, FDR 0.21), gene set. n = biologically independent mouse samples. GSEA was considered statistically significant for FDR (q-value) < 25% (**D**, **G**).

**Expanded View Figure 6.**
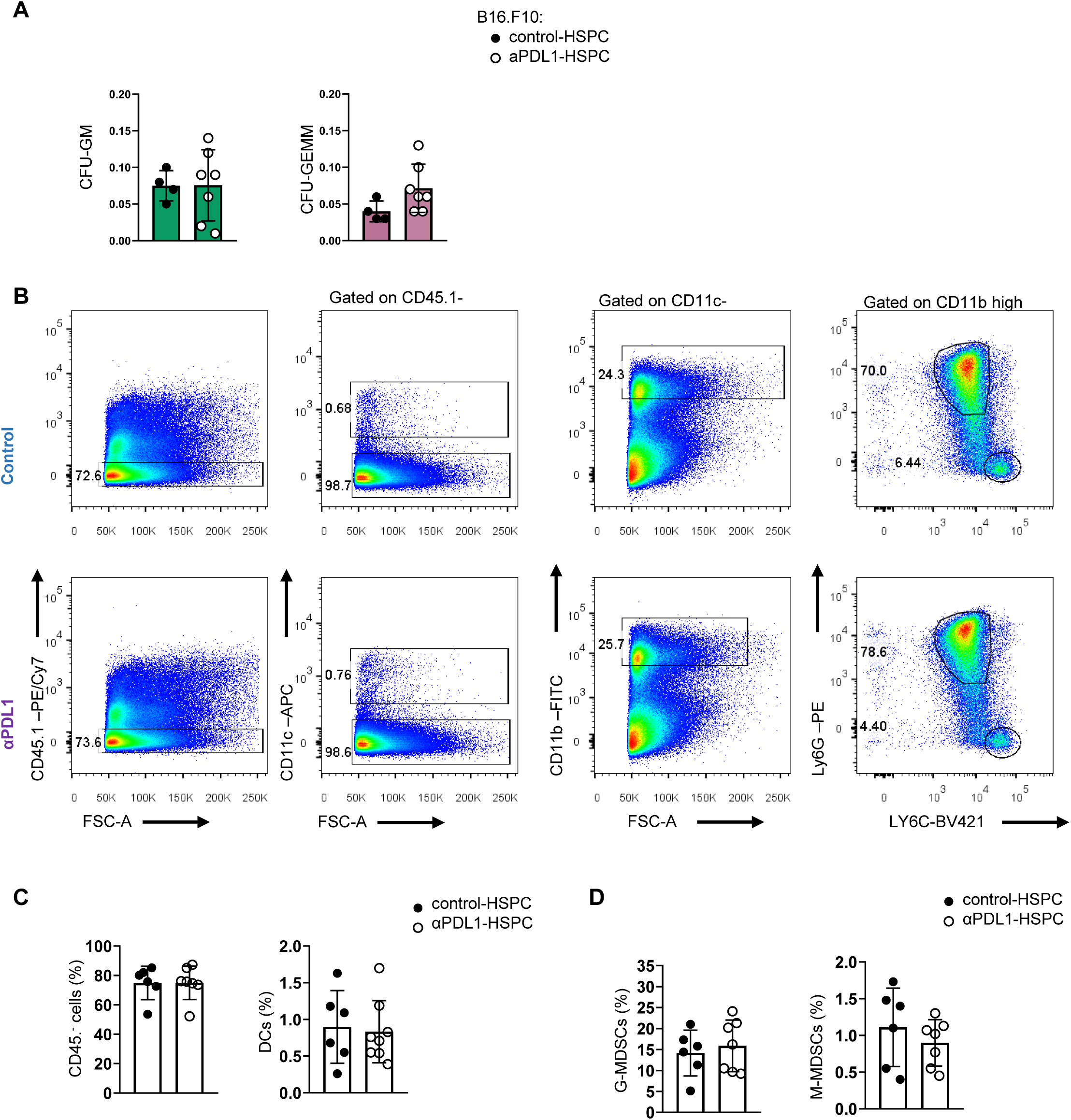
Successful engraftment and myelopoiesis followed transplantation of aPDL1-treated HSPCs in donor mice. (**A**) Normalized frequencies of CFU-GEMM, and CFU-GM of the PBS (control-HSPC, n = 4) and αPDL1 (αPDL1-HSPC, n = 7) conditions from 4 combined independent experiments (as in Figure 6B). (**B**) Gating strategy of the splenic CD45.1^-^, DCs cells, G-MDSCs, and M-MDSCs isolated from naive NBSGW mice transplanted with isolated HSPC (as in Figure6**C**; 1). Numbers denote percentages of gated populations. (**C**-**D**) Frequencies of splenic CD45.1^-^ cells, DCs cells (**C**; n = 6 control-HSPC, n = 7 αPDL1-HSPC) and in the CD11c^-^ population G-MDSCs, and M-MDSCs (**D**; n = 6 control-HSPC, n = 7 αPDL1-HSPC) isolated from naïve NBSGW mice (as in Figure 6C; 1). Representative data from two combined independent experiments out the five independent that were performed. p < 0.05*, p < 0.01**, p < 0.001***, p < 0.0001****. Means and SEM are depicted in all bar plots. If not stated otherwise, unpaired two-tailed t-tests are performed. N = biologically independent mouse samples

## Notes

The authors declare no potential conflicts of interest.

